# Apolipoprotein E controls Dectin-1-dependent development of monocyte-derived alveolar macrophages upon pulmonary β-glucan-induced inflammatory adaptation

**DOI:** 10.1101/2022.09.15.505390

**Authors:** H. Theobald, D.A. Bejarano, N. Katzmarski, J. Haub, J. Schulte-Schrepping, J. Yu, K Bassler, B. Ćirović, C. Osei-Sarpong, F. Piattini, L Vornholz, X. Yu, S. Sheoran, A. Al Jawazneh, S. Chakarov, K Haendler, G.D. Brown, D.L. Williams, L. Bosurgi, F. Ginhoux, J. Ruland, M. Beyer, M. Greter, M. Kopf, J.L. Schultze, A. Schlitzer

## Abstract

The lung is constantly exposed to the outside world and optimal adaptation of immune responses is crucial for efficient pathogen clearance. However, mechanisms which lead to the functional and developmental adaptation of lung-associated macrophages remain elusive. To reveal such mechanisms, we developed a reductionist model of environmental intranasal β-glucan exposure, allowing for the detailed interrogation of molecular mechanisms of pulmonal macrophage adaptation. Employing single-cell transcriptomics, high dimensional imaging and flow cytometric characterization paired to *in vivo* and *ex vivo* challenge models, we reveal that pulmonary low-grade inflammation results in the development of Dectin-1 - Card9 signaling-dependent monocyte-derived macrophages (MoAM). MoAMs expressed high levels of CD11b, ApoE, Gpnmb and Ccl6, were glycolytic and produced large amounts of interleukin 6 upon restimulation. Myeloid cell specific ApoE ablation inhibited monocyte to MoAM differentiation dependent on M-CSF secretion, promoting MoAM cell death thus impeding MoAM maintenance. *In vivo*, β-glucan-elicited MoAMs limited the bacterial burden of *Legionella pneumophilia* post infection and ameliorated fibrosis severity in a murine fibrosis model. Collectively these data identify MoAMs that are generated upon environmental cues and ApoE as an important determinant for lung immune resilience.

## Introduction

The lung is the body’s largest surface to interface with the outside world and is exposed to a variety of immunostimulatory agents shaping the lung’s ability to mount immune responses (Kopf et al., 2014). However, how such environmental non-clinically apparent immune activation is controlled on the cellular and molecular level is poorly understood. β-glucans are integral components of environmental pathogenic and non-pathogenic fungi and have been proposed as major systemic immune modulators (Divangahi et al., 2021). Furthermore, ambient concentrations of β-glucan, have been shown to oscillate over the course of the year and in combination with pathogen exposure have been correlated to an increase in allergic rhinitis (Shah and Panjabi, 2014). The recognition of β-glucan by Dectin-1 is crucial to modulate systemic immune responses through a process termed innate immune memory (Quintin et al., 2012), characterized by increased cytokine responses of trained innate immune cells, such as monocytes, upon a secondary non-related stimulus, facilitated by a metabolic and epigenetic rewiring, allowing a more efficient first line immune response (Arts et al., 2016a; Arts et al., 2016b; Bekkering et al., 2018; Cirovic et al., 2020; Quintin *et al.,* 2012; Saeed et al., 2014). However, the mechanisms by which β-glucan asserts its actions on the lung, the organ where it is most often recognized, and how it acts as a non-genetic modifier of immune response to subsequent disease, remain unknown.

The immune cell compartment of the - alveolar space of the lung is the body’s respiratory first line of defence, ensuring efficient immune response induction against airborne pathogens, while regulating immune activation to ensure intact lung function (Hussell and Bell, 2014). Within the - alveolar space alveolar macrophages (AM) and monocytes constitute the major mononuclear phagocytes found during homeostasis in humans and mice. Both AMs and monocytes express high amounts of Dectin-1 and exert highly plastic immune responses within the - alveolar compartment (Brown, 2005). Dectin-1 signalling has been shown to depend critically on spleen tyrosine kinase (Syk) which upon activation can trigger phospholipase C gamma 2 (PLCγ2) dependent calcium release and downstream nuclear factor of activated T-cells (NFAT) and extracellular signal-regulated kinase (ERK) activation. This signalling cascade leads to production of interleukin (IL) 2 and 10. Furthermore Syk can activate CARD9 leading to proinflammatory nuclear factor kappa-light-chain-enhancer of activated B cells (NFκB) signalling and the release of tumor necrosis factor alpha (TNFα) and IL-6. Additionally, Dectin-1 ligation has been described to directly induce reactive oxygen species (ROS) production via the activation of phosphoinositide 3 kinase (PI3K) (Gross et al., 2006; Vornholz and Ruland, 2020). Murine homeostatic AMs are embryonically derived with only minor contribution of adult bone marrow homeostasis (Guilliams et al., 2013; Schneider et al., 2014). However, viral infection or radiation of the lung have been shown to induce differentiation of Ly6c^+^ monocytes into long-lived monocyte-derived macrophages (Aegerter et al., 2020; Gibbings et al., 2017; Machiels et al., 2017). During viral infection monocyte-derived AMs have been shown to be crucial for the defence against subsequent *Streptococcus pneumoniae* infections through increased release of IL-6 (Aegerter *et al.,* 2020). Finally, environmental adaptation through viral exposure was also shown to affect the interaction between resident AMs and CD8^+^ T-cells, allowing a more efficient bacterial clearance of the latter upon viral training (Yao et al., 2018). Chronic lung diseases, such as asthma, idiopathic lung fibrosis or chronic obstructive pulmonary disease can be driven and accelerated by environmental factors. Therefore, it is crucial to understand the cellular, functional and molecular consequences of the immune recognition of ambient stimuli, such as β-glucan, and how the homeostatic broncho-alveolar resident mononuclear phagocyte compartment adapts in origin and function (Kaur et al., 2015; Lloyd and Marsland, 2017; Murdoch and Lloyd, 2010; Rabe and Watz, 2017).

Therefore, to investigate the impact of a non-pathological sterile environmental stimulus in a controlled manner we developed a reductionist model of a single low dose intranasal β-glucan exposure. Using single-cell transcriptomic, *in vivo* and *ex vivo* functional analysis of cellular development and downstream immune responses, this model allowed us to monitor acute and long-term effects on the broncho-alveolar mononuclear phagocyte compartment, and to dissect the molecular adaptation of macrophages upon environmental cues. We show that a single intranasal exposure to β-glucan induces the development of functionally modified monocyte-derived Apolipoprotein E (ApoE)^+^CD1 1b^+^ AMs, which are detected up to 21 days post β-glucan exposure. ApoE^+^CD11b^+^ AMs are glycolytic and release, upon restimulation with lipopolysaccharide (LPS), high amounts of IL-6. Furthermore, prior environmental adaptation by β-glucan followed by infection with *Legionella pneumophila* or a challenge by bleomycin-induced fibrosis led to an improved outcome after the secondary challenges. Molecularly, development of β-glucan induced ApoE^+^CD11b^+^ AM is controlled by the Dectin-1 - Card9 signalling pathway whereas maintenance of ApoE^+^CD11b^+^ AMs depends on paracrine ApoE and macrophage colony stimulating factor (M-CSF). Taken together, we identify the Dectin-1 - Card9 and ApoE pathways as critical regulators of functional environmental adaption of pulmonary mononuclear phagocytes and reveal the molecular constituents of such adaption processes, allowing potential future pharmaceutical intervention.

## Results

### Intranasal β-glucan exposure generates environmentally adapted ApoE^+^CD11b^+^ alveolar macrophages within the broncho-alveolar space

The lung is constantly exposed to the outside world, including pollutants, sterile and non-sterile pathogens and components thereof. However, knowledge about cellular and molecular mechanisms contributing to immune adaptation towards such environmental cues remains scarce. To reveal such mechanisms, we designed a reductionist model of a single low-dose particulate intranasal β-glucan exposure (200 μg) emulating environmental exposure (Rathnayake et al., 2017). To analyse the impact of this stimulation and to probe the longevity of its impact we assayed the macrophage compartment in the broncho-alveolar lavage fluid (BALF) (CD45^+^Lin^−^SSC^int-hi^) of PBS or β-glucan treated C57BL/6 mice seven days after inoculation using single-cell transcriptomics (**Figure 1A, S1A-C**). Dimensionality reduction by uniform-manifold approximation and projection analysis (UMAP) and unsupervised clustering using the Louvain algorithm revealed a total of five transcriptionally distinct clusters in the murine broncho-alveolar compartment (**Figure 1A, B**). High expression of the lung macrophage signature genes *Siglecf* and *Itgax* demonstrated that all investigated cells were of the macrophage lineage (**Figure S1D, E**) (Yu et al., 2017). Further analysis identified Clusters 0 and 1 as two subsets of resident AMs expressing *Ear2, Wfdc21* and *Fabp4, Hmox1,* respectively (Gautier et al., 2012; Yu *et al.*, 2017). Cluster 2 presented with expression of genes associated to proliferative processes such as *Top2a*, *Mki67* and *Birc5* and was therefore termed proliferating AMs (Cohen et al., 2018). Cluster 3 (ApoE^+^ AM) expressed many genes previously associated to a lipid-associated inflammatory monocyte-derived macrophage phenotype such as *Apoe, Cd63, Spp1, Gpnmb* and *Trem2* (Aegerter *et al.,* 2020; Jaitin et al., 2019; Keren-Shaul et al., 2017). Lastly, cluster 4 (ISG^+^ AM) was marked by expression of interferon-stimulated genes (ISG), such as *Ifit2, Ifit3, Ifi204* and *Isg15* (**Figure 1B**). Next, to understand whether one of the identified clusters was associated with β-glucan-induced environmental adaptation, we analysed the relative contribution of each condition to the clusters (**Figure 1C-E, S1F**). This revealed that ApoE^+^ AMs were only found within the broncho-alveolar space of mice exposed to β-glucan seven days prior, alongside a reduction of proliferating AMs. Previous work indicated CD11b expression on AMs post stimulation as a marker of enhanced inflammatory potential (Halstead et al., 2018; McCubbrey et al., 2018). Therefore, we utilized co-detection by indexing (CODEX) enabled high-dimensional imaging (Goltsev et al., 2018) to further describe the protein-level phenotype of ApoE^+^ AMs in the lung seven days post β-glucan stimulation (**Figure 1F**). This revealed that ApoE^+^ AMs co-expressed classical AM markers, such as CD11c and SiglecF, but also expressed significant amounts of CD11b, ApoE and GPNMB (**Figure 1F**). Therefore, we term this AM subpopulation ApoE^+^CD11b^+^ AMs. To further investigate the cellular dynamics of ApoE^+^CD11b^+^ AMs upon β-glucan induced environmental adaptation we monitored the BALF 0-21 days post β-glucan exposure using flow cytometry (**Figure 1G, S1G, H**). Flow cytometric analysis of the broncho-alveolar compartment showed that, in alignment with the single-cell transcriptomic data (**Figure 1E**), ApoE^+^CD11b^+^ AMs peaked at day seven post β-glucan inoculation and slowly diminished until day 21 (**Figure 1G**). Furthermore, in alignment with these data we found an overall increase in total AMs peaking at day seven post β-glucan exposure (**Figure S1H**). Next we verified ApoE^+^CD11b^+^ AMs within complete lung single cell suspension alongside their upregulated level of Siglec F, confirming their AM identity (**Figure S1I, J**).

**Figure 1.**
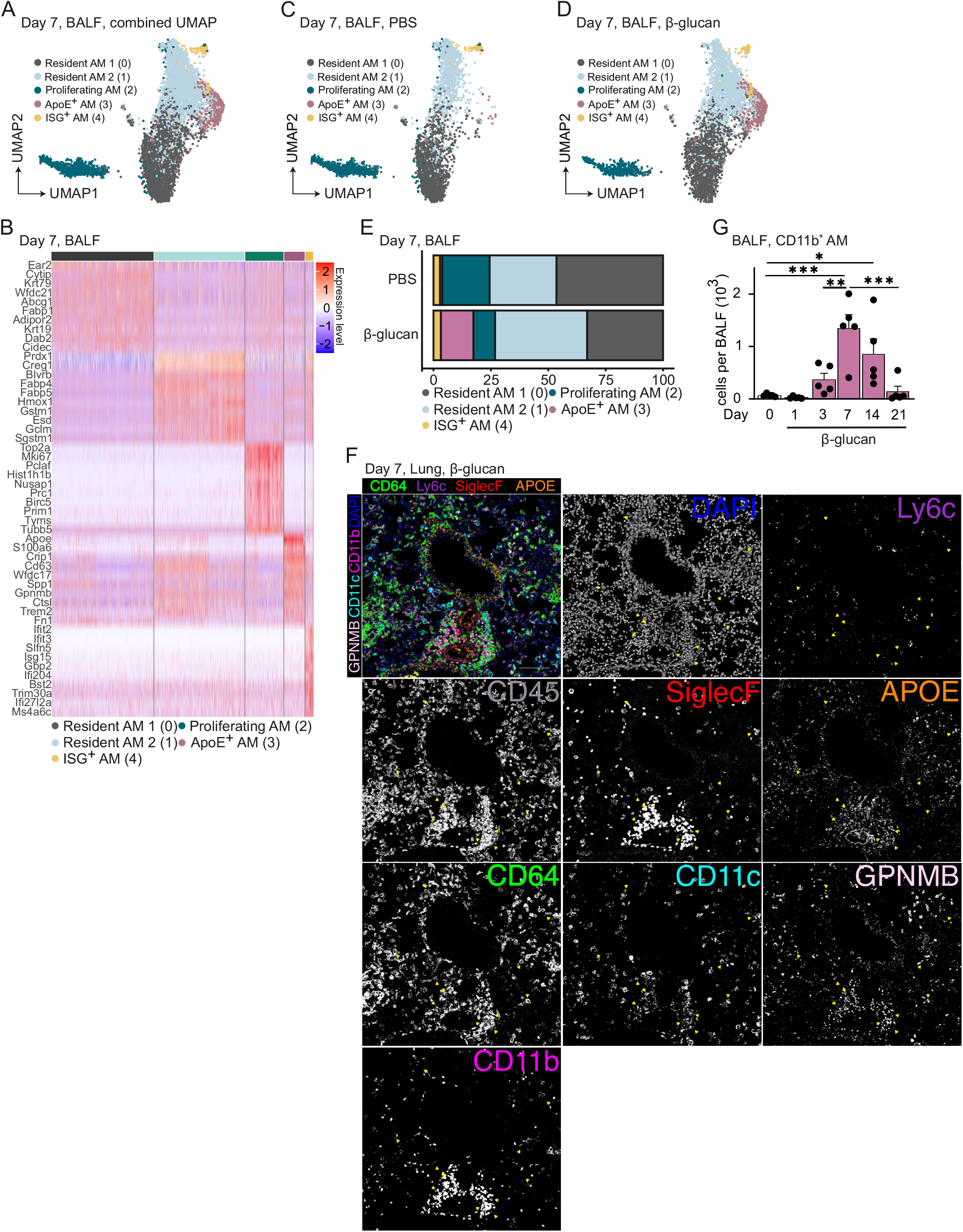
Intranasal β-glucan exposure generates environmentally adapted ApoE^+^CD11b^+^ alveolar macrophages within the broncho-alveolar space. (A-E) Single-cell RNA sequencing of the broncho-alveolar lavage fluid (BALF) of male 8-12 weeks old C57BL/6JCrl mice after intranasal stimulation with 40 μl 200 μg β-glucan or PBS (n=10202 cells). Seven days after exposure, cells were harvested and sorted for SSC^hi^, Lin^−^ (B220, CD19, CD3ε, Nk1.1, Ter-119), DRAQ7^−^ singlets. (A) Uniform-manifold approximation and projection (UMAP) analysis of both conditions combined shows five different clusters. (B) Heatmap of top ten highly expressed genes for each of the five clusters. (C, D) UMAP from (A) separated by PBS (C, n= 4845 cells) or β-glucan (D, n= 5357 cells) condition. (E) Percentage of contribution of the five annotated clusters to overall cells split by conditions. (F) 5μm frozen section of the left lobe of the lung of a *Ms4a3-cre^Rosa26TOMATO^* mouse 7 days after β-glucan exposure stained with a 17-plex CODEX antibody panel. Overlay image shows the markers used to identify AM populations. Single stainings of these markers are shown in grayscale. Arrows indicate CD11b^+^ ApoE^+^ AMs. Scale bar represents 100μm. (G) Flow cytometric quantification of absolute ApoE^+^CD11b^+^ AM (CD45^+^SiglecF^+^CD64^+^CD11c^+^CD11b^+^) numbers in the BALF in a time course from 1 to 21 days post β-glucan stimulation of wt mice (n= 5).

### ApoE^+^CD11b^+^ AMs are monocyte-derived and CCR2-dependent

Previous data indicated that acute and chronic inflammation can lead to the recruitment of monocyte-derived macrophages into the broncho-alveolar space (Aegerter *et al.,* 2020; McCubbrey *et al.,* 2018; Mould et al., 2017). To understand whether β-glucan induced environmental adaptation rewires a resident alveolar macrophages subpopulation or leads to recruitment of Ly6c^+^ monocytes and their subsequent differentiation into ApoE^+^CD11b^+^ AMs we first monitored the influx of Ly6c^+^ monocytes in the broncho-alveolar space using flow cytometry. This revealed that post β-glucan stimulation Ly6c^+^ monocytes were recruited to the broncho-alveolar space peaking on day three after stimulation, remaining high ad day 7 and decreasing from day 14 onwards (**Figure 2A**). To connect this data with the emergence of ApoE^+^CD11b^+^ AMs we employed the *Ms4a3-cre^Rosa26TOMATO^* mouse model which allows genetic lineage tracing of the bone marrow derived monocyte lineage. Genetic lineage tracing revealed that 71% ± 10% ApoE^+^CD11b^+^ AMs were labelled with tomato indicating an overall bone marrow monocyte origin (**Figure 2B, C**). Additionally, CODEX imaging revealed co-expression of CD11b and dtTomato in SiglecF^+^CD11c^+^ AMs within the lung, further adding weight to their monocyte origin (**Figure 2D**). To further validate this finding we employed the CCR2^−/−^ mouse model in which recruitment of Ly6c^+^ monocytes upon inflammation is strongly decreased (Serbina and Pamer, 2006). To determine whether generation of environmental adaptation induced ApoE^+^CD11b^+^ AMs is CCR2 dependent, we inoculated either control or CCR2^−/−^ mice with β-glucan and analysed the BALF seven days later using flow cytometry (**Figure 2E**). This analysis revealed a significant reduction of ApoE^+^CD11b^+^ AMs upon β-glucan inoculation in CCR2^−/−^ mice. Taken together these results demonstrate that β-glucan-induced ApoE^+^CD11b^+^ AMs arise from bone marrow progenitors in a CCR2-dependent fashion.

**Figure 2.**
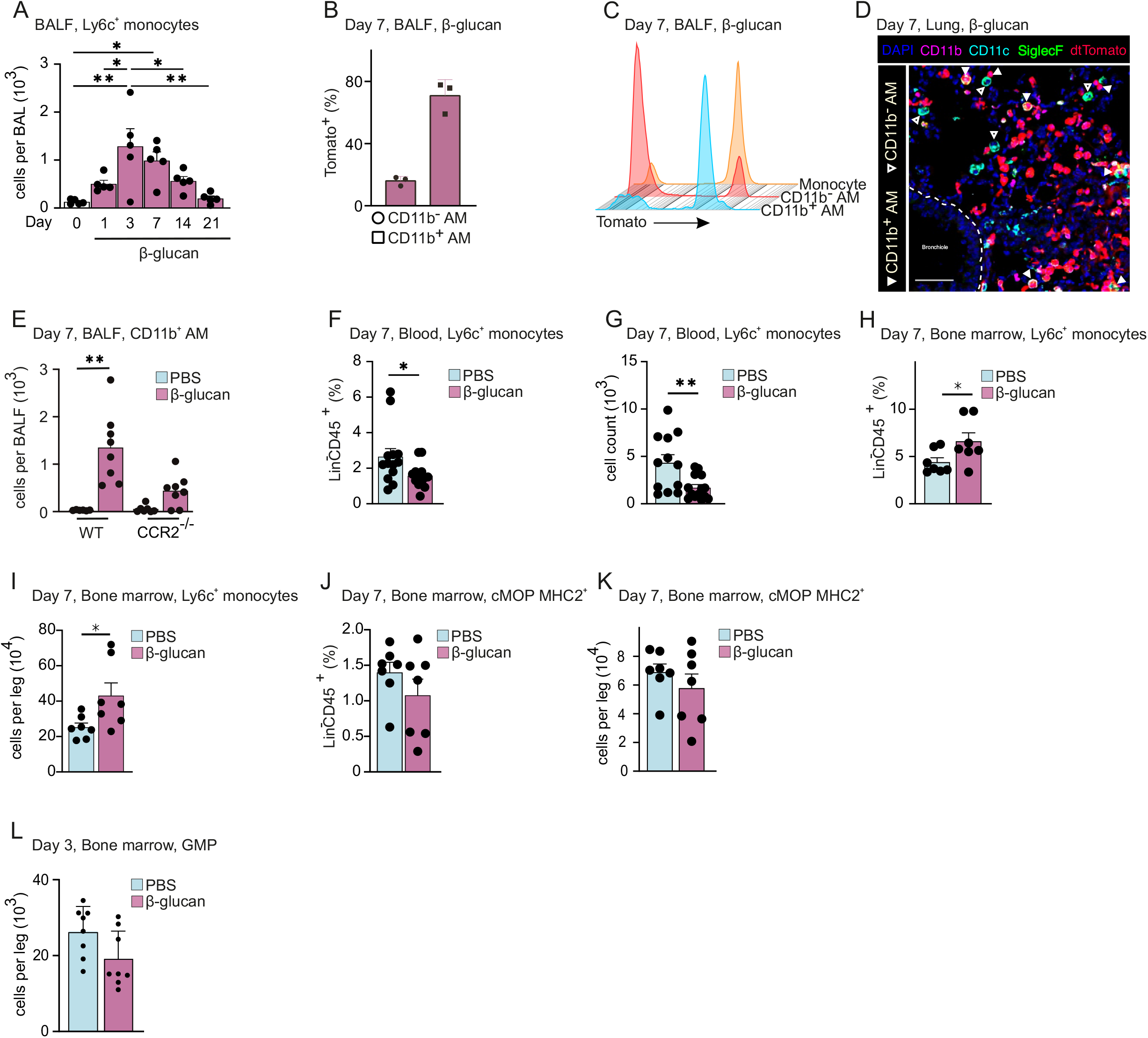
ApoE^+^CD11b^+^ AMs are monocyte-derived and CCR2-dependent. (A) Flow cytometric quantification of absolute Ly6c^+^ monocyte (CD45^+^Ly6g^−^SiglecF^−^CD64^+^CD11b^+^Ly6c^+^) numbers in the BALF 1 to 21 days post β-glucan stimulation of C57BL/6J mice (n=4-5). (B, C) Flow cytometric analysis of BALF from *Ms4a3-cre^Rosa26TOMATO^* mice seven days after intranasal PBS or β-glucan stimulation (n=3). (B) Percentage of Tomato^+^ labelling in CD11b^−^ and CD11b^+^ AM and (C) proportion of Tomato^+^ labelling in CD11b^−^ and CD11b^+^ AM compared to monocytes (CD45^+^SiglecF^−^Ly6g^−^CD11b^+^F4/80^+^). (D) CODEX multiplexed immunostaining of the left lobe of a *Ms4a3-cre^Rosa26TOMATO^* mouse 7 days after β-glucan exposure (enlargement from Fig. 1F). Filled arrowheads indicate CD11b^+^ ApoE^+^ AMs, whereas empty arrowheads indicate CD11b^−^ AMs. dtTomato reporter signals are represented in red. Scale bar represents 50μm. (E) Flow cytometric quantification of absolute CD11b^+^ AM numbers in the BALF seven days after β-glucan exposure in wt or CCR2^−/−^ mice (n=8). (F-K) Percentage and absolute counts of Ly6c^+^ monocytes in the blood (F, G, n=12) or Ly6c^+^ monocytes and MHCII^+^ cMOPs in the bone marrow (H-K, n=4) of C57BL/6J mice seven days after PBS or β-glucan by flow cytometry. (K) Flow cytometric quantification of GMPs in the bone marrow of C57BL/6J mice three days after intranasal PBS or β-glucan stimulation by flow cytometry (n = 8).

### β-glucan-elicited environmental adaptation increases Ly6c^+^ monocyte differentiation within the bone marrow

Systemic β-glucan administration can lead to changes in abundance and functional priming of myeloid-associated precursors within the bone marrow (Kalafati et al., 2020; Mitroulis et al., 2018). To reveal whether or not local broncho-alveolar adaptation elicits systemic feedback mechanisms affecting bone marrow monopoiesis, we analysed abundance of Ly6c^+^ monocytes in blood and bone marrow seven days post intranasal β-glucan stimulation using flow cytometry. Here, we found a significant reduction of both percentage and absolute cell count of Ly6c^+^ monocytes in blood and an increased percentage and number of Ly6c^+^ monocytes within the bone marrow (**Figure 2F-I**). To understand if the increased number of bone marrow Ly6c^+^ monocytes is a function of an increased precursor reservoir, we analysed the bone marrow resident common monocyte progenitor (cMOP) repertoire of mice stimulated intranasally with β-glucan or PBS seven days before. This revealed that activated cMOPs, expressing MHC2, were decreased in mice pre-treated with β-glucan (**Figure 2J, K**). Thereby these data indicate that enhanced generation of Ly6c^+^ monocytes upon intranasal β-glucan mediated environmental adaptation evokes MHC2^+^ cMOP-mediated differentiation to allow generation of sufficient peripheral Ly6c^+^ monocytes. Furthermore, we addressed whether enhanced peripheral demand for Ly6c^+^ monocytes and their progenitors affect the abundance of granulocyte macrophage progenitors (GMP) in the bone marrow of mice intranasally stimulated with β-glucan or PBS. Here, we found a trend towards reduced GMP numbers in the bone marrow of mice seven days after treatment with β-glucan compared to PBS (**Figure 2L**). Taken together, these data indicate a peripheral feedback mechanism which draws upon immediate monocyte progenitors to fill up demand of monocytes within the lung, without significantly affecting the myeloid differentiation cascade *per se*.

### ApoE^+^CD11b^+^ AMs show increased release of interleukin 6 and induction of glycolysis

Monocyte-derived macrophages have been associated with enhanced inflammatory response after strong inflammatory and infectious events, such as influenza A infection (Aegerter *et al.,* 2020). However, if and how the mononuclear cell repertoire of the broncho-alveolar space functionally adapts to low grade inflammatory stimuli, remains unclear. To investigate whether BALF-resident macrophages exhibit a different functional profile seven days post β-glucan exposure, we isolated BALF-resident immune cells, enriched for macrophages by adherence, and re-stimulated macrophages with PBS or LPS for 24h *in vitro*. Next, supernatants were investigated for IL-6 and TNFα release using ELISA. This revealed that macrophages pre-treated with β-glucan released significantly higher amounts of IL-6 and a trend towards more TNFα after LPS re-stimulation as compared to PBS pre-treated with PBS (**Figure 3A, S2A**). To further validate that indeed ApoE^+^CD11b^+^ AMs were endowed with a higher capacity to release IL-6 post β-glucan stimulation, we utilized intracellular flow cytometric analysis of IL-6 secretion after LPS or PBS restimulation. Here, we found a strong increase in IL-6^+^CD11b^+^ AMs post β-glucan exposure as compared to the PBS exposed controls (**Figure 3B, C, S2B, C**). Our data shows that generation of ApoE^+^CD11b^+^ AMs upon β-glucan exposure is CCR2-dependent (**Figure 2C**). Therefore, to validate that the increased IL-6 within *ex vivo* restimulated BALF resident macrophage cultures can be attributed directly to ApoE^+^CD11b^+^ AMs, we exposed CCR2^−/−^ and control mice to β-glucan and seven days later enriched macrophages from the BALF by adherence and restimulated adherent cells with LPS for 24hs *in vitro* (**Figure 3D**). This analysis revealed that the increased amount of IL-6 seen in β-glucan-exposed broncho-alveolar macrophages is CCR2-dependent, showing that ApoE^+^CD11b^+^ macrophages are the major source of increased IL-6 upon β-glucan-induced environmental adaptation. Furthermore, enhanced cytokine responses upon systemic β-glucan stimulation have been linked to the induction of glycolysis within mononuclear phagocytes (Arts *et al.*, 2016b; Bekkering *et al.*, 2018). To understand whether this was also true in our system, we utilized extracellular flux analysis to measure glycolysis. This analysis showed that glycolysis, glycolytic capacity and reserve were significantly increased on day seven post β-glucan exposure in adherence-selected BALF-resident macrophages (**Figure 3E, F, S2D, E**). Collectively, these data indicate an enhanced proinflammatory potential of ApoE^+^CD11b^+^ AMs.

**Figure 3.**
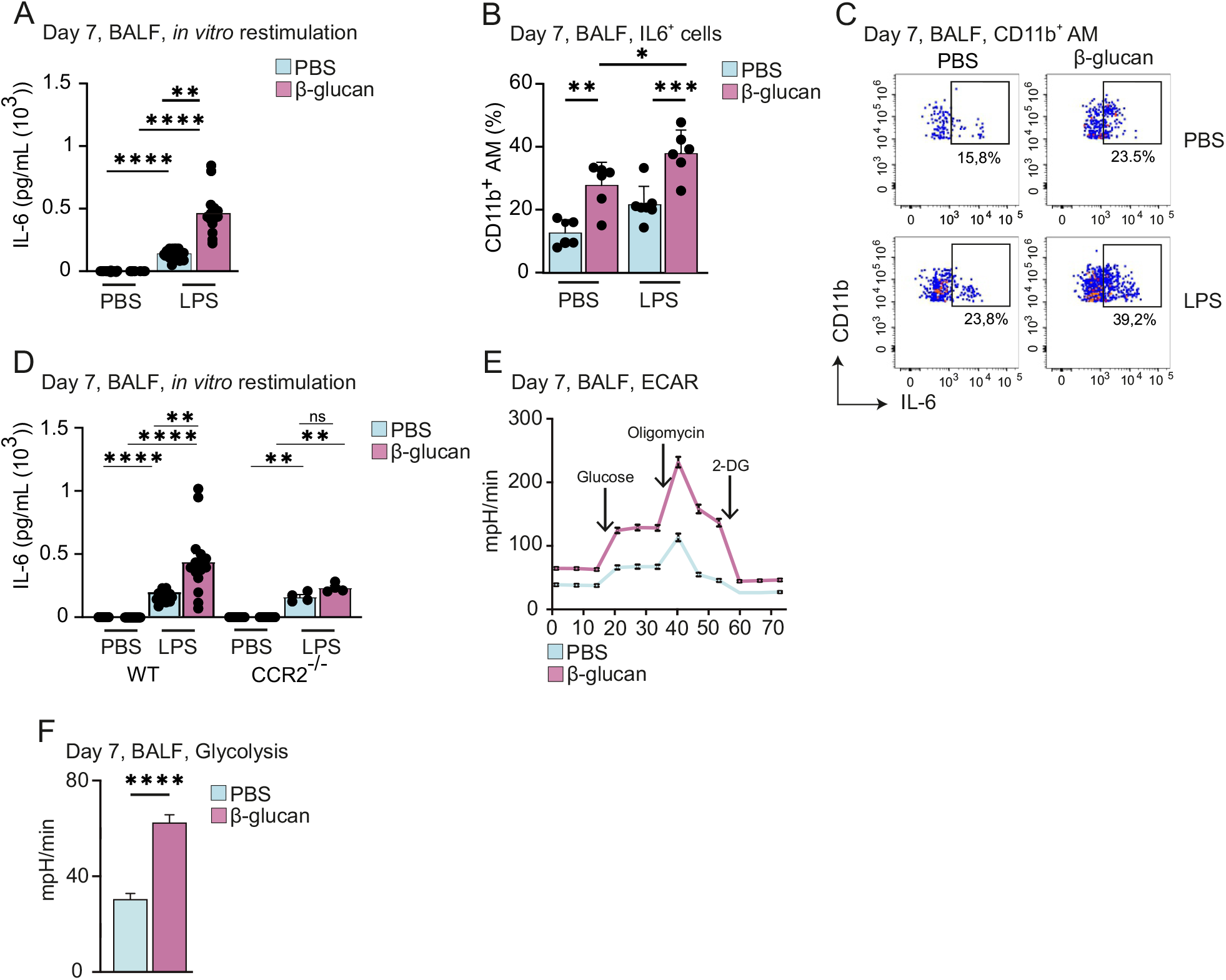
ApoE^+^CD11b^+^ AMs show increased release of interleukin 6 and induction of glycolysis. (A-C) BALF cells were harvested from C57BL/6J mice seven days after intranasal exposure with PBS or β-glucan and subsequently restimulated *in vitro* with PBS or LPS. (A) Quantification of IL-6 protein levels by ELISA in the cell culture supernatant 24 h after LPS restimulation (n=15). (B) Intracellular staining and flow cytometric measurement of IL-6. Dot plot shows the percentage of IL-6^+^ CD11b^+^ AM after restimulation with LPS for 6-8 h followed by intracellular staining and flow cytometric analysis (n=6). (C) Representative flow cytometry dot plots of (B) gated on CD11b^+^ AM. (D) Quantification of IL-6 protein levels by ELISA in the cell culture supernatant 24 h after restimulation with LPS of C57BL/6J (n=15) and CCR2^−/−^ mice (n=4). (E, F) Extracellular flux analysis acidification rate (E) and glycolysis (F) in BALF cells seven days after PBS or β-glucan stimulation (n=6).

### β-glucan-induced environmental adaptation enhances pulmonary bacterial clearance and ameliorates bleomycin induced fibrosis

Optimal adaptation to outside stimuli received via ambient air is essential for a potent, yet measured immune response. IL-6 has been indicated to be a major component of an adapted pulmonary immune response. As ApoE^+^CD11b^+^ AMs produced increased amounts of IL-6, we wondered whether those cells have an impact on the outcome of an acute and chronic inflammation. To dissect whether this enhanced cytokine secretion ability affects protection against a bacterial pathogen, we utilized C57B/L6 mice, which were environmentally adapted for seven days, infected these mice intratracheally with *Legionella pneumophila* and analysed the bacterial burden and cellular composition of the BALF two days post infection **(Figure S3A)**. Here, environmentally adapted mice showed a significant decrease of bacteria detected within their BALFs and an increased count of proinflammatory macrophages associated to bacterial clearance (**Figure 4A, B**). Next, to investigate if environmental adaptation has consequences beyond the modulation of acute bacterial infection, we inoculated environmentally adapted or control mice intratracheally with bleomycin to induce experimental lung fibrosis (**Figure S3B**). Here, environmentally adapted mice showed a significantly higher rate of survival over a 14-day observation period post bleomycin inoculation alongside a trend towards a reduced area of lung fibrosis as examined by histology (**Figure 4C, D-E**). Supporting this notion, environmentally adapted mice also showed a lower score of disease burden and a decreased loss of bodyweight during lung fibrosis (**Figure S3C, D**). Furthermore, molecular effectors associated to fibrosis resolution, such as IL-4 and IL-33 were enhanced early after bleomycin inoculation (day 3), whereas profibrotic TSLP (day 14) was decreased during the fibrotic phase in environmentally adapted mice (**Figure S3E-G**). Collectively these functional data reveal a profound regulatory role of environmental adaption for the regulation and severity of acute and chronic inflammation, likely regulated by ApoE^+^CD11b^+^ AMs.

**Figure 4.**
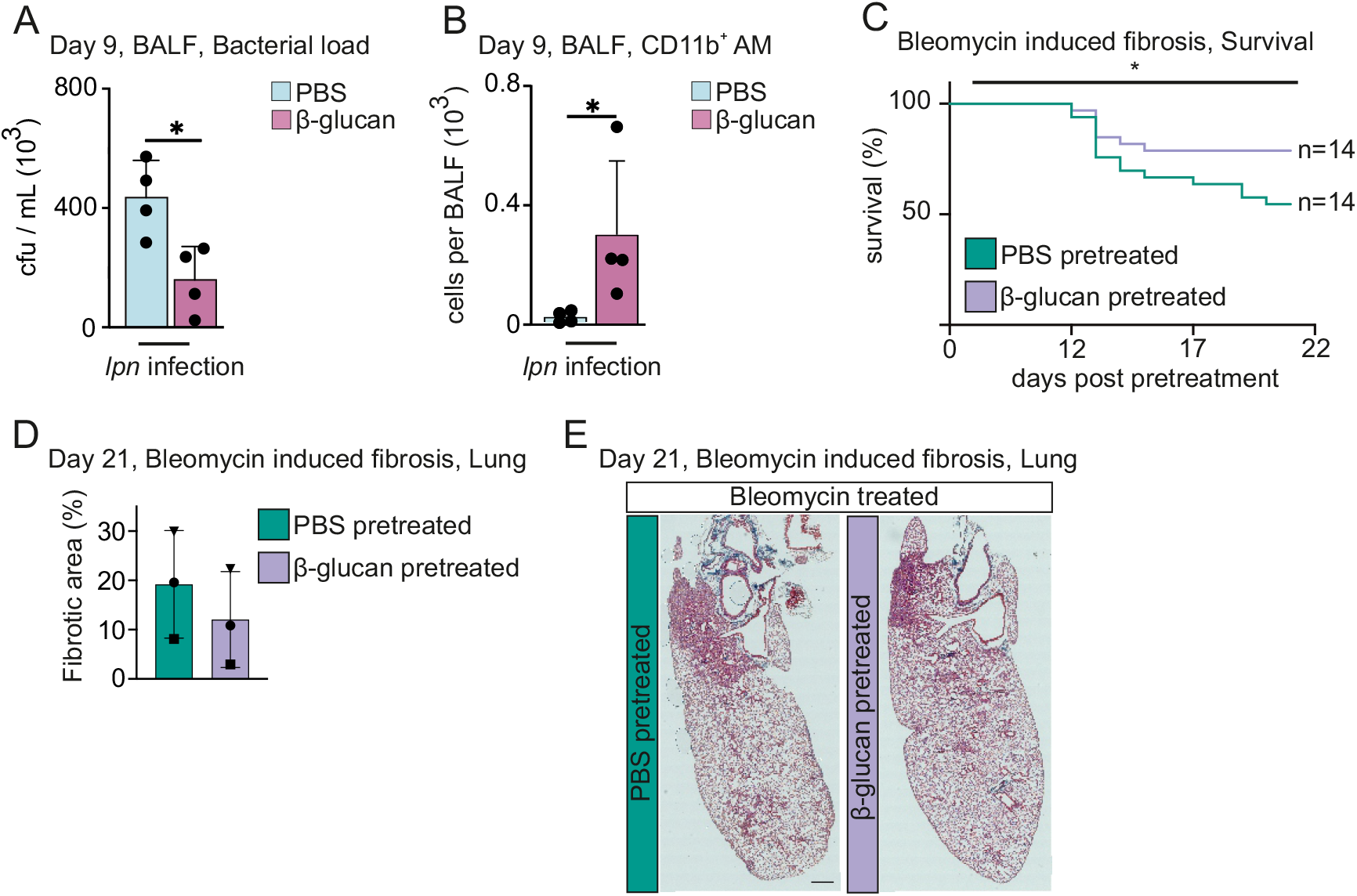
β-glucan-induced environmental adaptation enhances pulmonary bacterial clearance and ameliorates bleomycin-induced fibrosis. (A, B) C57BL/6J mice were intranasally stimulated with PBS or β-glucan followed by intratracheal infection with *Legionella pneumophilia* at day seven post primary stimulation and analysis at day nine (n=4). Quantification of bacterial load in BALF (A) and absolute numbers of CD11b^+^ AM by flow cytometry nine days post primary stimulation (B). (C-E) Pulmonary fibrosis was induced seven days after PBS or β-glucan stimulation by intratracheal gavage of *Streptomyces verticillus* bleomycin. (C) Mice were monitored daily for survival (n=14). (D-E) Representative images of Masson’s Trichome staining of lung tissue at day 21 after initial stimulation (E) and quantification of fibrotic areas (D). Each dot in (D) represents a mouse. Scale bar in (E) represents 500 μm.

### Generation of ApoE^+^ CD11b^+^ AMs by β-glucan is dependent on the Dectin-1 – Card9 signalling axis

β-glucan is recognized by various receptors, including CR3, Dectin-1 and CD5 (Fesel and Zuccaro, 2016; Kalia et al., 2021; Taylor et al., 2007). The most prominently expressed receptor on mononuclear phagocytes is Dectin-1. To further understand how development of ApoE^+^CD11b^+^ AMs is regulated on the molecular level, we profiled expression of Dectin-1 on BALF-resident macrophages using flow cytometry (**Figure 5A**). This revealed that homeostatic Dectin-1 expression is largely confined to the resident AM compartment with only a small fraction of monocytes expressing Dectin-1. Further to assess whether expression of Dectin-1 is necessary for the development of ApoE^+^CD11b^+^ AMs, we intranasally inoculated either control or Dectin1^−/−^ mice with β-glucan and analysed the cellular composition of BALF resident immune cells seven days later using flow cytometry. This analysis revealed that generation of ApoE^+^CD11b^+^ AMs is dependent on Dectin-1 expression, whereas initial inflammatory recruitment of Ly6c^+^ monocytes to the air-exposed space is not (**Figure 5B, C**). Next, to understand whether immune cell-intrinsic or stromal cell recognition via Dectin-1 is critical for the development of ApoE^+^CD11b^+^ AMs and thus environmental adaptation, we transferred Dectin1 ^−/−^ or control (CD45.2^+^) bone marrow into lethally irradiated CD45.1^+^ control mice and analysed their BALF compartment seven days post environmental adaption by β-glucan. Here, generation of ApoE^+^CD11b^+^ AMs was entirely dependent on hematopoietic expression of Dectin-1 (**Figure 5D**). CARD9 has been shown to be a critical molecule for Dectin-1-mediated Nfκb activation (Deerhake et al., 2021; Gross *et al.,* 2006; Vornholz and Ruland, 2020). To investigate whether or not ApoE^+^CD11b^+^ AMs require Card9 for Dectin-1 signalling in order to develop, we treated lethally irradiated mice reconstituted with Card9^−/−^ or control BM with β-glucan or solvent and analysed the BALF seven days later by flow cytometry. This revealed that development of ApoE^+^CD11b^+^ AMs depends on Dectin-1-elicited Card9-dependent signalling pathways (**Figure 5E**). Next, we investigated whether the loss of ApoE^+^CD11b^+^ AMs by abrogating Dectin-1 or Card9 signalling also leads to a loss of increased IL-6 secretion upon *in vitro* restimulation with LPS in adherence-selected macrophages. In line with the data obtained in the CCR2^−/−^ mouse model, enhanced IL-6 secretion was abolished in the absence of Dectin-1 or Card9 signalling and thus can be attributed to ApoE^+^CD11b^+^ AMs (**Figure 5F, G**).

**Figure 5.**
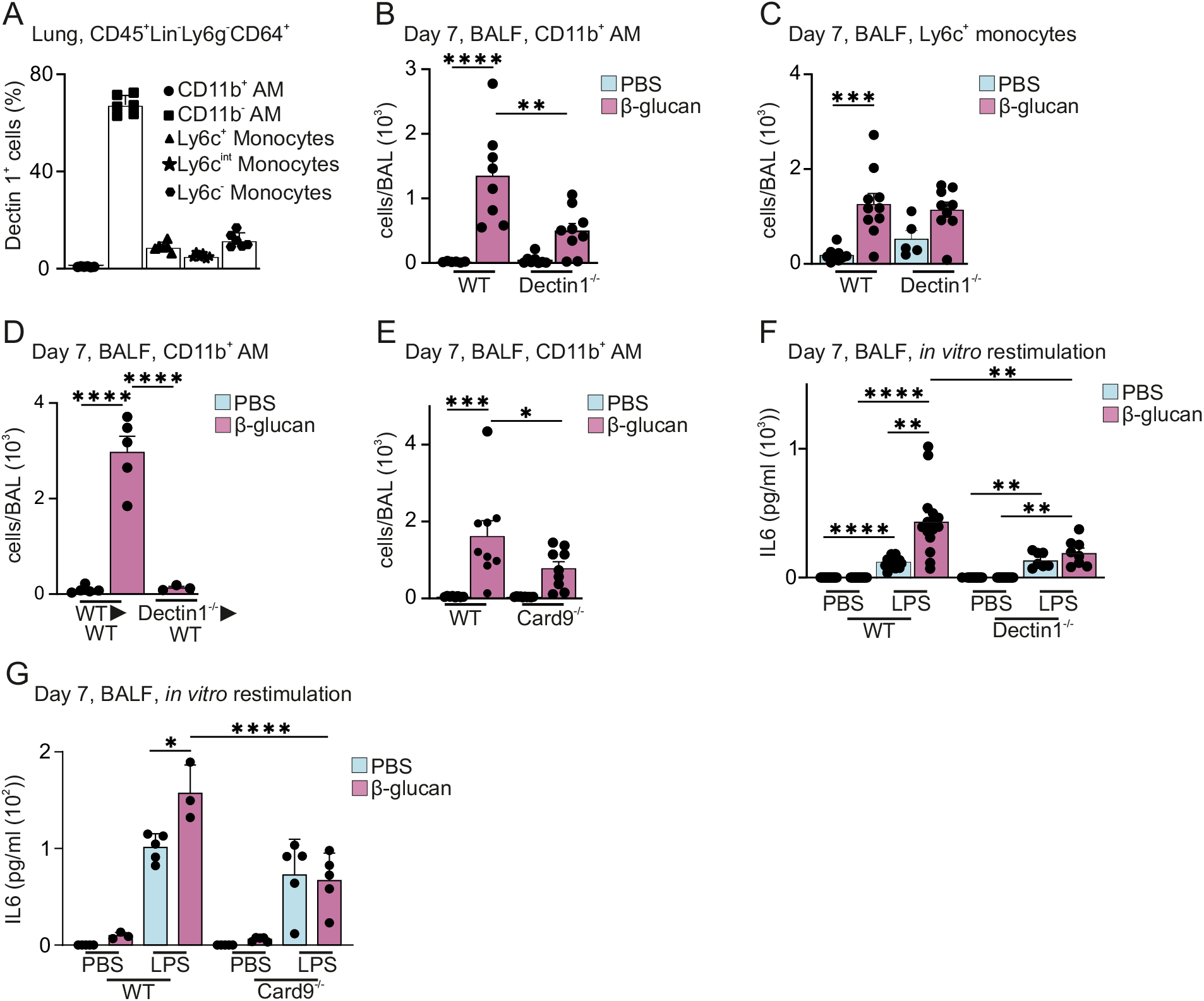
Generation of ApoE^+^CD11b^+^ AMs by β-glucan is dependent on the Dectin-1 – Card9 signalling axis. (A) Percentage of monocyte and macrophage populations contributing to Dectin-1^+^ cells in the C57BL/6J mouse lung pregated on CD45^+^Lin^−^Ly6g^−^CD64^+^ cells (n=6) by flow cytometry. (B, C) Absolute CD11b^+^ AM (B) and Ly6c^+^ monocyte (C) numbers in the BALF seven days after PBS or β-glucan exposure in C57BL/6J or Dectin1^−/−^ mice (n=5-9) by flow cytometry. (D, E) Absolute CD11b^+^ AM numbers in the BALF seven days after PBS or β-glucan exposure in Dectin1^−/−^ (D, n=3-6) or Card9^−/−^ (E, n=8-9) bone marrow chimeras by flow cytometry. (F) Quantification of IL-6 protein levels by ELISA in the cell culture supernatant 24 h after LPS restimulation of C57BL/6J (n=15) and Dectin1^−/−^ mice (n=7-8). (G) Quantification of IL-6 protein levels by ELISA in the cell culture supernatant 24 h after LPS restimulation of C57BL/6J (n=3-5) and Card9^−/−^ mice (n=5).

### Paracrine myeloid-derived Apolipoprotein E controls ApoE^+^CD11b^+^ AM differentiation upon β-glucan-induced environmental adaptation

*Apoe* is highly expressed across a range of monocyte-derived macrophage populations identified in various different low grade or chronic inflammatory diseases, however its exact role in the monocyte to macrophage differentiation and maintenance has not been studied (Aegerter et al., 2020; Jaitin et al., 2019). Similarly, in our model of β-glucan-induced environmental adaptation, *Apoe* was abundantly expressed in CD11b^+^ AMs and was also readily detectable on the protein level as early as one day post intranasal β-glucan stimulation within the BALF (**Figure 6A, B**). To further clarify the role of ApoE for environmental adaptation of the mononuclear phagocyte repertoire in the lung, we intranasally inoculated ApoE^flox^Lysm^cre^ mice with β-glucan, which specifically lack ApoE expression within the myeloid lineage, and analysed the composition of the BALF-resident mononuclear phagocyte compartment seven days later using flow cytometry. Here, β-glucan stimulated ApoE^flox^Lysm^cre^ mice failed to generate CD11b^+^ AMs and in line with this did not have increased numbers of Ly6c^+^ monocytes within their BALF (**Figure 6C, D**). Next, we wondered whether ApoE was controlling the generation of ApoE^+^CD11b^+^ AMs in a paracrine or autocrine manner as both signalling modes have been described before (Dove et al., 2005; Kemp et al., 2021). Therefore we generated mixed (50:50) wildtype (CD45.1^+^) : ApoE^flox^Lysm^cre^ (CD45.2^+^) bone marrow chimeras or congenic mixed wildtype (CD45.1^+^) : wildtype (CD45.2^+^) control chimeras. After reconstitution, chimeras were stimulated intranasally with β-glucan and their BALF-resident mononuclear phagocyte repertoire was analysed using flow cytometry seven days later. This analysis revealed that both WT/WT and WT/ApoE^flox^Lysm^cre^ chimeras efficiently generated ApoE^+^CD11b^+^ AMs seven days post β-glucan exposure supporting a paracrine signalling mode (**Figure 6E**, **S4A**). Next, we assessed the contribution of ApoE-deficient CD45.2^+^ cells to the pool of ApoE^+^CD11b^+^ AMs. This revealed that both ApoE-proficient (CD45.1) and deficient (CD45.2) cells equally contributed to the pool of ApoE^+^CD11b^+^ AMs post β-glucan stimulation (**Figure 6F**). These data reveal that a paracrine myeloid cell derived source of ApoE is sufficient to rescue the generation of ApoE^+^CD11b^+^ AMs upon environmental adaptation in the lung.

**Figure 6.**
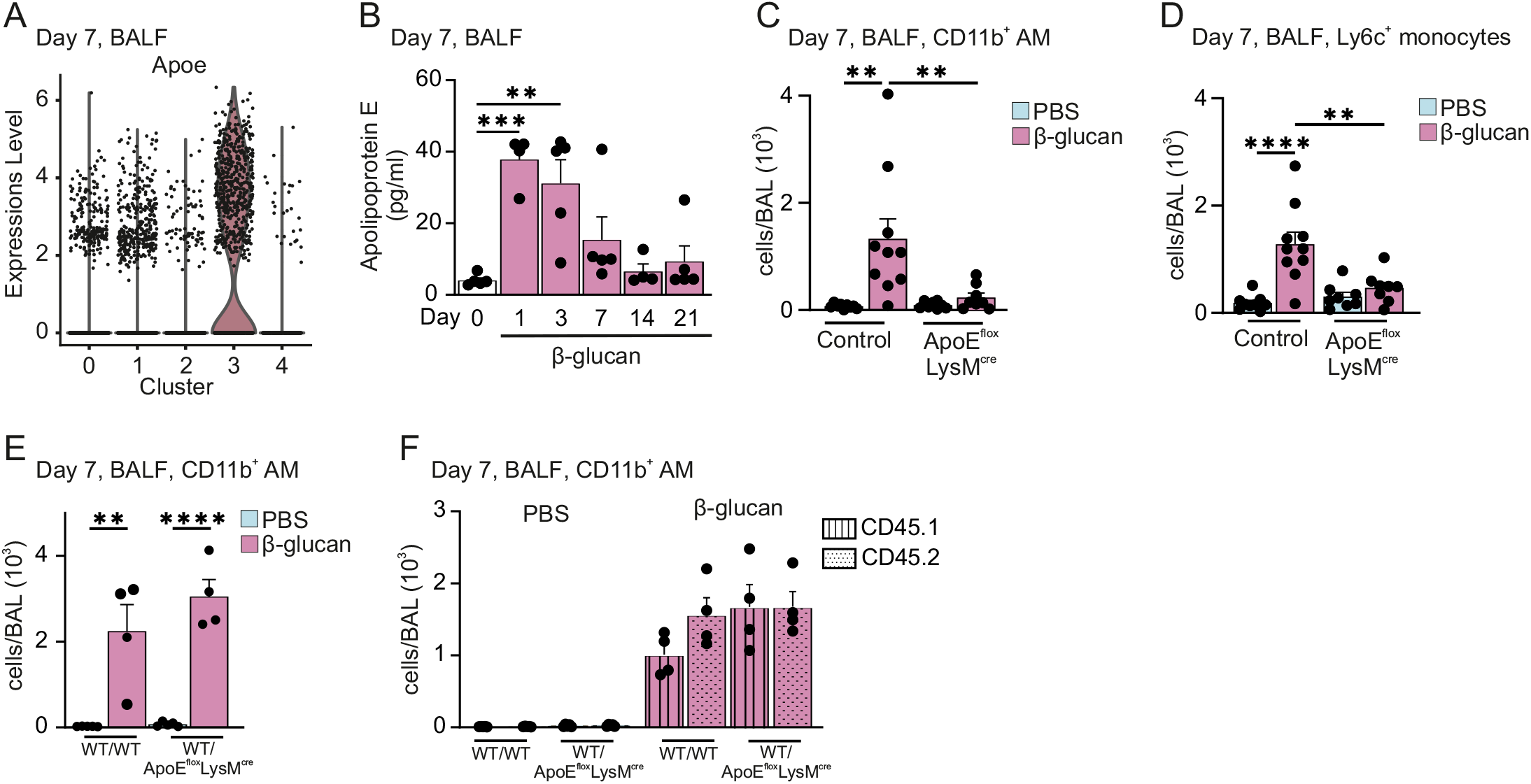
Paracrine myeloid-derived Apolipoprotein E controls ApoE^+^CD11b^+^ AM differentiation upon β-glucan-induced environmental adaptation. (A) Violin plot of *ApoE* RNA expression levels in the BALF seven days after β-glucan exposure by scRNA seq. (B) C57BL/6J mice were stimulated with β-glucan and BALF was harvested at different time points. Plot shows ApoE protein levels in the BALF measured by ELISA (n=5). (C, D) Absolute numbers of CD11b^+^ AM (C) and Ly6c^+^ monocytes (D) seven days after intranasal β-glucan exposure of C57BL/6J or ApoE^flox^LysM^cre^ mice by flow cytometry (n=8-10). (E, F) Lethally irradiated CD45.1^+^/CD45.2^+^ male mice were reconstituted with 1.5×10^6^ CD45.1^+^ mixed with CD45.2^+^ bone marrow cells (WT/WT) or with CD45.1^+^ mixed with ApoE^flox^LysM^cre^ CD45.2^+^ bone marrow cells (WT/ ApoE^flox^LysM^cre^) for 12 weeks and subsequently intranasally stimulated with PBS or β-glucan (n=4-5). Flow cytometric quantification of CD11b^+^ AM numbers (E) and contribution of donor cells (CD45.1^+^ or CD45.2^+^) to the CD11b^+^ AM pool (F) seven days after exposure.

### Myeloid-derived Apolipoprotein E controls survival of ApoE^+^CD11b^+^ AMs by regulation of cholesterol storage and M-CSF secretion

To reveal the potential mechanism of action for ApoE in the differentiation process of ApoE^+^CD11b^+^ AMs, we investigated whether myeloid-derived ApoE is crucial for the initial commitment of Ly6c^+^ monocytes towards monocyte-derived macrophage development or if maintenance or survival of monocyte-derived macrophages is controlled by ApoE. To study this, we analysed wildtype or ApoE^flox^Lysm^cre^ mice three days post intranasal inoculation with β-glucan. Here, ApoE^+^CD11b^+^ AMs and Ly6c^+^ monocyte numbers within the BALF were comparable between control and ApoE^flox^Lysm^cre^ mice indicating that initial commitment towards the macrophage lineage and recruitment of monocytic precursors to air-exposed space is not controlled by ApoE, but establishing a critical time window of action for ApoE between day three and day seven post β-glucan inoculation (**Figure 7A, B**). To determine whether ApoE regulates survival of ApoE^+^CD11b^+^ AMs and thus their subsequent longevity *in vivo*, we assessed ApoE^+^CD11b^+^ AM survival three days post β-glucan challenge using TUNEL staining. This analysis showed increased abundance of TUNEL^+^ApoE^+^CD1 1b^+^ AMs in the absence of myeloid derived ApoE upon β-glucan challenge indicating a possible role of ApoE in the regulation of ApoE^+^CD11b^+^ AM survival (**Figure 7C**). To further reveal the possible molecular dysregulation caused by the lack of ApoE in myeloid cells, we evaluated intracellular cholesterol content of BALF resident CD11b^+^ AMs three days post β-glucan stimulation. This revealed that upon β-glucan stimulation ApoE-deficient CD11b^+^ AMs showed an increase in intracellular cholesterol content as indicated by filipin staining (**Figure 7D, E**). Increased intracellular cholesterol storage has been associated to cholesterol induced toxicity by dysregulation of intracellular protein transport (Song et al., 2021). Furthermore, it was shown that autocrine production of M-CSF is crucial for the BALF-resident survival of CD11b^+^ AMs upon viral infection (Wu et al., 2015). To investigate whether dysregulated M-CSF production due to cholesterol accumulation is involved, we quantified the amount of M-CSF within the BALF of mice intranasally stimulated with β-glucan for 24h. These data showed production of M-CSF within the BALF 24h post β-glucan stimulation in dependence of intact myeloid cell derived ApoE signalling (**Figure 7F, S5A**). Furthermore, fluorescent microscopy revealed that the majority of M-CSF^+^ cells within the lung also expressed Siglec F and CD11b (**Figure S5B, C**). To understand whether the observed decrease of soluble M-CSF can be attributed to ApoE^+^CD11b^+^ AMs we isolated BALF-resident macrophages by adherence from either control or ApoE^flox^Lysm^cre^ mice and assessed the production of M-CSF by CD11b^+^ AMs by microscopy (**Figure 7G**). This showed that ApoE-proficient CD11b^+^ AMs produced increased amounts of M-CSF 24h after β-glucan stimulation whereas ApoE-deficient CD11b^+^ AMs lost their ability to produce M-CSF in response to β-glucan stimulation. Taken together this suggests ApoE as a central regulator of monocyte to macrophage differentiation and survival via the M-CSF signalling axis within the lung upon β-glucan-induced environmental adaptation.

**Figure 7.**
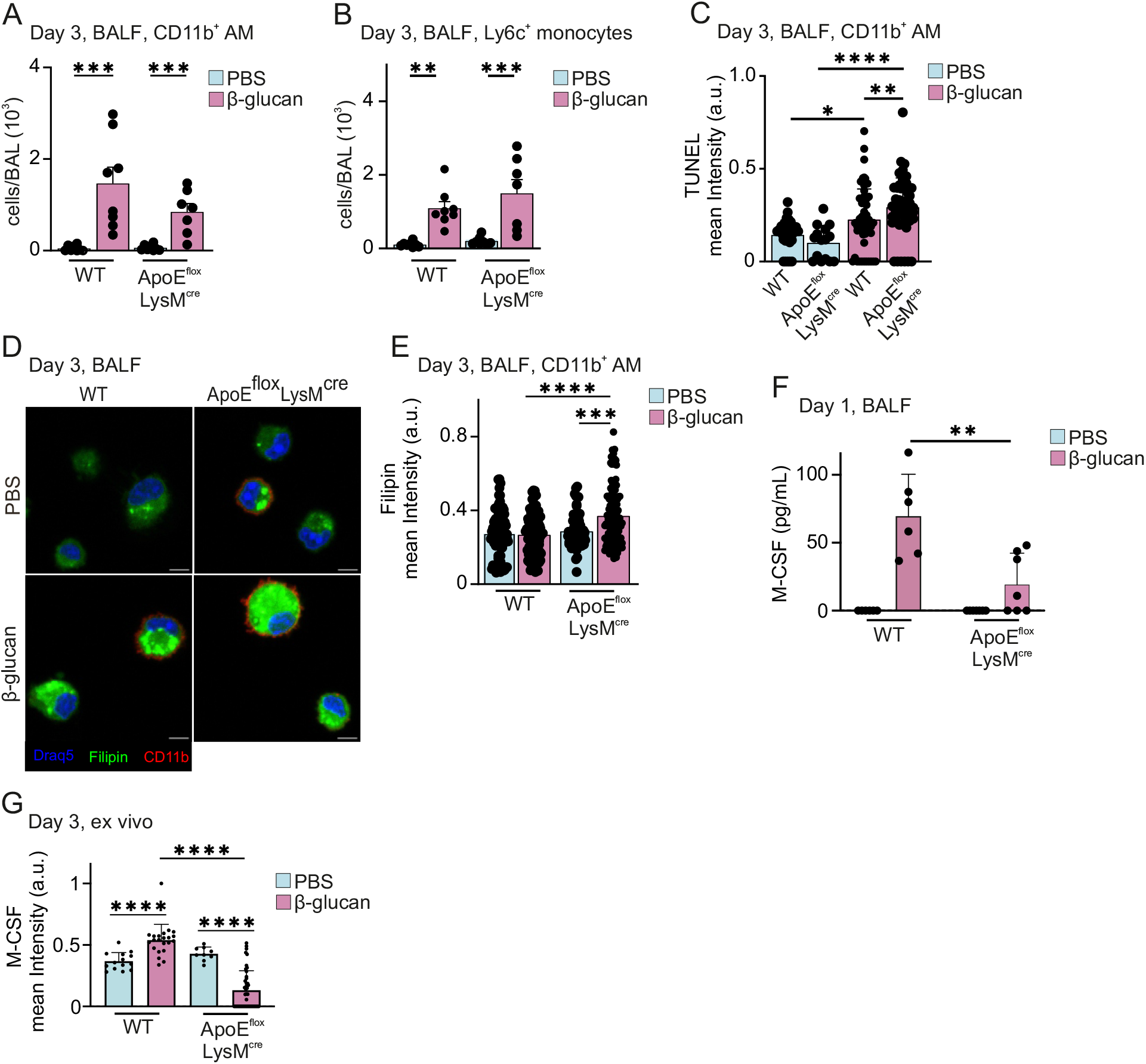
Myeloid-derived Apolipoprotein E controls survival of ApoE^+^CD11b^+^ AMs by regulation of cholesterol storage and M-CSF secretion. (A-B) Absolute numbers of CD11b^+^ AM (A) and Ly6c^+^ monocytes (B) three days after intranasal β-glucan exposure in C57BL/6J or ApoE^flox^LysM^cre^ mice (n=7-8) by flow cytometry. (C-E) C57BL/6J and ApoE^flox^LysM^cre^ mice were stimulated with PBS or β-glucan intranasally. Three days after stimulation, BALF was harvested, seeded and fixed after 2 h. (C) TUNEL staining was performed, followed by conventional immunofluorescence to detect SiglecF and CD11b. Plot shows mean TUNEL signal intensities of individual CD11b^+^ AM (n=4). (D, E) Filipin staining after fixation and immunofluorescence analysis. Representative images of filipin stainings are shown in (D). Scale bars represent 5 μm. Plot in (E) shows mean filipin signal intensities of individual CD11b^+^ AM in the different conditions (n=5). (F) Quantification of M-CSF protein levels in the BALF one day after β-glucan exposure in C57BL/6J mice or ApoE^flox^LysM^cre^ mice (n=6) measured by ELISA. (G) C57BL/6J and ApoE^flox^LysM^cre^ mice were stimulated with PBS or β-glucan intranasally. 24 h after stimulation BALF was harvested and seeded. Cells were fixed after 2h and immunostained to detect SiglecF, CD11b, and M-CSF. Plot shows mean M-CSF signal intensities of individual CD11b^+^ AMs (n=4).

## Discussion

Within our modern-day environment, the lung is constantly exposed to a plethora of sterile immunostimulatory components. However, the immunological and molecular consequences for lung-resident macrophage development and function are incompletely understood. Here, we show that a single non-pathologic intranasal β-glucan stimulus induces the development of monocyte-derived alveolar macrophages, which highly express CD11b and ApoE and are further characterised by their superior IL-6 production capacity in response to secondary LPS stimulation. Additionally, CD11b^+^ApoE^+^ AMs are glycolytic and modify the outcome of a secondary bacterial infection and of a chronic fibrotic response *in vivo.* Molecularly this adaptation is instructed via the recognition of β-glucan by Dectin-1 and signal transduction via its adaptor protein CARD9. Further examination of the molecular requirements of macrophage adaptation revealed a crucial role of ApoE for maintenance of CD11b^+^ApoE^+^ AMs within the broncho-alveolar environment via the control of macrophage-derived M-CSF. These data reveal that ApoE is a crucial determinant for environmental macrophage adaptation and couples cellular cholesterol metabolism to cellular environmental adaptation.

Earlier studies have examined the development of monocyte-derived AMs during pathological insults, such as viral infections, radiation or bleomycin-induced fibrosis (Aegerter *et al.,* 2020; Li et al., 2022a; Machiels *et al.,* 2017; Misharin et al., 2017). However, how broncho-alveolar macrophages adapt their transcriptome, metabolism and function to ambient immune stimulatory components, such as LPS or β-glucan, after the acute or sub-acute recognition phase remains poorly understood (Zahalka et al., 2022). Here, we show that although the initial inflammation evoked by β-glucan is minimal, functionally modified monocyte-derived AMs arise from Ly6c^+^ monocytes within the air-exposed space of the lung, a process affiliated to strong pathology-induction, as e.g. found during viral infection. This validates the model as a suitable for low grade environmentally induced inflammation. CD11b^+^ApoE^+^ AMs are induced for up to 21 days post β-glucan stimulation leading to functional modification of the macrophage repertoire in the lung, demonstrating the importance of low-grade inflammatory sterile insults in shaping the overall immune competence of the lung-resident macrophage repertoire.

β-glucan is an integral cell wall component of many pathological and non-pathological fungi and can be found within ambient air (Rathnayake *et al.,* 2017). Recent studies evaluated the role of β-glucan for the induction of innate immune training on the systemic level but failed to examine its effects at the level of the tissues exposed (Novakovic et al., 2016; Quintin *et al.,* 2012). Here, β-glucan-induced functional modulations were accompanied by the induction of glycolysis and the enhanced release of the pro-inflammatory cytokines IL-6 and TNFα in circulating Ly6^+^ monocytes. These findings are in line with our results of the β-glucan-induced functional adaptation in the lung. Furthermore, systemic β-glucan administration was reported to modify bone marrow myeloid development, by expansion of specific GMP / MPP like progenitors, leading to outcomes ranging from enhance pathogen clearance to maladaptation (Kalafati *et al.,* 2020; Li et al., 2022b; Mitroulis *et al.,* 2018). In line with these findings, we show that topical subclinical stimulation of the lung with β-glucan only minimally affects bone marrow resident progenitors and that generation of CD11b^+^ApoE^+^ AMs only depends on late stage direct monocytic progenitors and recruitment of Ly6c^+^ monocytes to the lung via CCR2-dependent signalling. Dectin-1 is crucial for the induction of inflammation upon β-glucan challenge, however its downstream signalling is heterogenous and determines the functional output. We show that tissue adaptation conveys Dectin-1 signalling via CARD9-dependent signalling circuits, in concordance with increased levels of IL-6, most probably via activation of Nfκb signalling.

Finally, we show that β-glucan-induced macrophages upregulate ApoE, a protein which has been demonstrated to be part of various disease-specific monocyte-derived macrophage gene signatures, e.g. during influenza infection, lung fibrosis or obesity (Aegerter *et al.,* 2020; Jaitin *et al.,* 2019; Misharin *et al.,* 2017). However, its functional role for monocyte-derived macrophage development was not investigated. In hematopoietic stem cells ApoE was shown to inhibit proliferation and subsequent progenitor maturation by controlling sensitivity towards granulocyte macrophage stimulating factor and IL-3 (Murphy et al., 2011). We show that myeloid-specific deletion of ApoE leads to accumulation of intracellular cholesterol, increased cell death and a reduction of autocrine M-CSF production ultimately inhibiting the formation of a long-lived monocyte-derived macrophage compartment upon intranasal β-glucan challenge. Generation of ApoE^+^ macrophages can also be found in white adipose tissues during obesity, a condition conferring training-like feature to mononuclear phagocytes or during influenza-induced lung inflammation supporting the crucial role of ApoE for the development of functionally adapted macrophages during inflammation (Christ et al., 2018; Jaitin *et al.,* 2019). Other work has established functional Nfκb-responsive elements within the ApoE reporter and CARD9 directly activates Nfκb thus directly couples activation of the inflammatory Nfκb response to the induction of inflammation experience in monocyte-derived macrophages.

Collectively we provide evidence that a single non-pathological environmental stimulation via the Dectin-1 – CARD9 axis generates inflammation-experienced monocyte-derived macrophages, which modify subsequent acute and chronic inflammation within the lung microenvironment under the control of ApoE, thus molecularly linking macrophage inflammatory amplitude to lung resilience and disease susceptibility.

## Acknowledgments

We thank the whole Schlitzer Lab for their fruitful discussion. This study was funded by the Deutsche Forschungsgemeinschaft (DFG, German Research Foundation) under Germany’s Excellence Strategy – EXC2151 – 390873048 (AS), and an Emmy Noether research grant (SCHL2116/1 to AS). This research was funded in part by Wellcome Trust [Grant number 217163 and 102705]. For the purpose of open access, the author has applied a CC BY public copyright licence to any Author Accepted Manuscript version arising from this submission. We also thank the Medical Research Council Centre for Medical Mycology at the University of Exeter for funding (MR/N006364/2). M.G. and X.X. were funded by the Swiss National Science Foundation (310030_184915). JLS is supported by the German Research foundation (DFG) under Germany’s Excellence Strategy (EXC2151-390873048) as well as under SCHU 950/8-1; GRK 2168, TP11; SFB704, the BMBF-funded excellence project Diet–Body–Brain (DietBB) and the EU project SYSCID under grant SFB 841number 733100.

## Methods

### Animal studies and mouse models

All mice used in this study were bred in the animal facility of the LIMES Institute, University of Bonn, Germany or Center for Translational Cancer Research, Klinikum rechts der Isar, Technical University of Munich, Germany. Animals were housed in IVC mice cages under conventional conditions (12 h/12 h light/dark cycle, 22°C), with ad libitum access to food and water. Eight to twelve weeks old male mice were used for experiments. The ApoE^flox^ mice were kindly provided by Prof. J. Heeren. All experiments were approved by government of North Rhine-Westphalia (84-02.04.2017.A347, 81-02.04.2020.A454).

### Intranasal stimulation

Mice were anesthetized by intraperitoneal injection of ketamine/xylazine and intranasally inoculated with endotoxin-free 1× PBS (EMD Millipore) or 200 μg β-glucan (⊕95% pure) from *Candida albicans*, which was provided by Prof. D. Williams. The broncho-alveolar lavage fluid (BALF) was collected one, three, seven, 14 or 21 days after inoculation. For cell analysis, the lung was flushed 3× with 1 ml cold 1× PBS with 10 mM EDTA. Afterwards, the fluid was centrifuged for 5 min at 1350 rpm at 4°C and the supernatant was discarded. For cytokine and chemokine assessment, the lung was flushed 3× with the same 1 ml of cold 1× PBS with 10 mM EDTA. Afterwards, the samples were centrifuged for 10 min at 14000 rpm at 4°C and the supernatant was frozen in liquid nitrogen until analysis. Supernatants were thawn on ice and centrifuged for 5 min at 10 000 rpm at 4°C to remove debris. M-CSF (R&D Systems) and ApoE (Abcam) protein levels were measured by ELISA according to manufacturer’s protocols.

### Pulmonary fibrosis and *Legionella pneumophila* infection

Mice were intranasally stimulated with β-glucan or PBS seven days before induction of pulmonary fibrosis or infection by *Legionella pneumophila*. For fibrosis induction, *Streptomyces verticillus* bleomycin (Sigma-Aldrich, 0.75 mg/kg body weight) was administered by intratracheal gavage. Body weight and health status were scored on a daily basis. Analysis was performed three or 14 days after bleomycin application. For bacterial infections, intratracheal application of *Legionella pneumophila* (5×10^6^ CFU/mouse) was performed and mice were sacrificed 2 days after induction. *L. pneumophila* was provided by A. Oxenius. Bacteria from the glycerol (25%) stock were plated on a CYE-plate and grown for 3 days at 37°C in a non-CO2 incubator. For bacterial load determination one lobe of the lung was collected in 1 mL PBS with 1 metal bead and smashed using a tissue lyser (Qiagen). 25 μL of lysed lung were put in duplicates on CYE-plates and grown for 3-4 days at 37°C in a non-CO_2_ incubator.

### Bone marrow Chimeras

Bone marrow chimeras were generated by multiple intraperitoneal busulfan (Sigma-Aldrich) injections or irradiation of recipient mice with 10 Gy. Afterwards, 5 × 10^6^ (busulfan-treated mice) or 1.5 ×10^6^ (irradiated mice) freshly isolated bone marrow cells from the donor animals were intravenously injected into the recipients. Peripheral blood chimerism was assessed 28 days after reconstitution by flow cytometry. Bone marrow chimeras were used in experiments after eight – twelve weeks of reconstitution.

### Flow cytometry and cell sorting

For flow cytometry, cells of the broncho-alveolar space were harvested by flushing the lungs with 3× 1 ml ice cold 1× PBS containing 10 mM EDTA. After centrifugation with 1350 rpm for 5 min at 4°C, cell pellets were resuspended in antibody mix and stained for 35 min at 4°C. After washing with FACS buffer (1× PBS, 2 mM EDTA, 0.5% BSA (SERVA)), life/death stain was performed using DRAQ7 (BioLegend, 1:1000 in FACS buffer) for 5 min at RT. Red blood cell lysis was performed only if necessary. In case of blood analysis, blood was collected in 1× PBS with 10 mM EDTA and stained in antibody mix for 35 min at 4°C. Red blood cell lysis was performed twice for 5 min at RT before life/death stain and acquisition. For bone marrow cells, femur and tibia were flushed with 1x PBS, stained with antibody mix for 1 h and red blood cell lysis and life/death stain were subsequently performed. Cells were washed and resuspended in FACS buffer and recorded using FACS Symphony A5 (Becton Dickinson). FACS data were analyzed using FlowJo v10.8.1 (Becton Dickinson).

Cell sorting was performed using an ARIA III (Becton Dickinson) instrument. Briefly, cells from the lung lavage were antibody stained followed by life/death stain. Cell sorting was performed using a 100 μm nozzle into cooled 1.5 ml reaction tubes containing FACS buffer.

### *Ex vivo* stimulation and assessment of cytokine production

BALF of stimulated mice was collected in 15 ml tubes, centrifuged at 1350 rpm for 5 min at 4°C and resuspended in 1 ml RPMI1640 (PAN Biotech) supplemented with 10% FCS (Sigma-Aldrich), 2 mM GlutaMAX (Gibco), 5 ml MEM non-essential amino acids (Sigma Aldrich), 1 mM Na-Pyruvate (Gibco), 50 U/ml Pen-Strep (Gibco) and 500 μl β-Mercaptoethanol. Cells were counted and seeded with 0.2 ×10^5^ cells per well. After 2 h resting in 500 μl medium at 37°C and 5% CO_2_, cells were stimulated with 10 ng/ml LPS (Sigma-Aldrich). After 24 h, cell culture supernatant was harvested and snap frozen for further analysis. Supernatants were thawn on ice and centrifuged for 5 min at 10 000 rpm at 4°C to remove debris. TNF-α and IL-6 (ThermoFisher) protein levels were measured by ELISA. For multiplex cytokine and chemokine analysis, a customized 18-plex Procartaplex kit (ThermoFisher) was used according to manufacturer’s protocols and run on Luminex FLEXMAP 3D (ThermoFisher) device.

### Intracellular cytokine stain

Lavage fluid of two mice was pooled and centrifuged for 5 min at 1350 rpm at 4°C. Cells were resuspended in 1 ml RPMI1640 (PAN Biotech) supplemented with 10% FCS (Sigma-Aldrich), 2 mM GlutaMAX (Gibco), 5 ml MEM non-essential amino acids (Sigma Aldrich), 1 mM Na-Pyruvate (Gibco), 50 U/ml Pen-Strep (Gibco) and 500 μl β-Mercaptoethanol. Equal cell numbers were plated into two wells per condition and rested in 500 μl media for 2 h at 37°C and 5% CO_2_. Cells were stimulated with 10 ng/ml LPS (Sigma-Aldrich) for 6 h in total. After 4 h, 2.5 μg Brefeldin A (BioLegend) and 2 nM Monensin (BioLegend) were added to each well and incubated for further 2 h. Afterwards, supernatant was discarded and cells were harvested in 1xPBS using a cell scraper followed by staining of surface markers by antibodies for 30 min at 4°C. Cells were washed and permeabilized using the Cytofix/Cytoperm kit (Becton Dickinson, adapted from manufacturer’s protocol). In brief, cells were resuspended in 200 μl Cytofix/Cytoperm solution per tube and incubated for 20 min at 4°C. Cells were washed twice with Perm/Wash and stained with Zombie NIR fixable viability dye (1:1000 in PBS) for 15 min. After washing with Perm/Wash, cells were intracellularly stained by 100 μl Perm/Wash containing IL-6 (MP5-20F3; 1:100) or TNF-α (MP6-XT22; 1:100) antibody or the corresponding isotype control for 30 min at 4°C. Cells were washed twice with Perm/Wash and resuspended in 1x PBS before acquisition.

### Seahorse analysis

BAL fluid of two mice was pooled and 50.000 – 100.000 cells were plated in a 96 well Seahorse plate (Agilent) in Seahorse XF base medium (Agilent) supplemented with 5% L-Glutamine (Sigma-Aldrich), 10% FCS (Sigma-Aldrich) and 50 U/ml Pen/Strep (Gibco) for 2 h at 37°C and 5% CO_2_. Before acquisition, cells were washed and incubated in Seahorse XF base medium with 5% L-Glutamine (Sigma-Aldrich) and 50 U/ml Pen/Strep (Gibco), but without FCS and Glucose for 30 min at 37°C without CO_2_ supply. During the Seahorse run, 100 mM Glucose (Sigma-Aldrich) solution was injected into port A leading to a final glucose concentration of 10 mM per well. This was followed by injection of 10 μM Oligomycin A (Sigma-Aldrich, final concentration 1 μm) solution and 500 mM 2-Desoxyglucose (Sigma-Aldrich, final concentration 50 mM). Glycolysis, glycolytic capacity and glycolytic reserve were calculated using the Agilent Wave software.

After the Seahorse assay, cell numbers of each well were determined using the CyQUANT NF Cell Quantification Assay (Thermo-Fisher) for normalization. 50 μl of CyQUANT NF Cell Quantification mix were added to each well, incubated for 30 min at RT and measured in a TECAN plate reader (excitation wavelength 485 nm, emission wavelength 530 nm).

### Histology

Mice were anesthetized followed by trans-cardial perfusion with 10ml ice cold 1xPBS containing 10 mM EDTA using the lung-heart circulation. Lungs were removed and fixed in 4% PFA overnight at 4°C (for paraffin-embedded tissue) or infiltrated with 1 mL 50% OCT (in 1×PBS), removed, and fixed for 6 h in 1.3% PFA at 4°C. For paraffin sections lungs were dehydrated and paraffin embedded. For frozen sections, after fixation lungs were dehydrated in 10, 20, and 30% sucrose (in 1× PBS) for 24 h at 4°C. After dehydration, the left lobe was separated and embedded in OCT. Sections of 5 μm were prepared for Hematoxylin and Eosin (H&E) and Trichrome (Masson) stainings and immunohistochemistry.

### Immunofluorescence and histology stainings

Coverslips containing frozen tissue sections were left drying on drierite beads for 5min and subsequently fixed on ice-cold acetone for 10 min. Afterwards, sections were washed twice with 0.01% Tween-PBS and permeabilized with 0.2% Triton X-100 for 20min at RT. After permeabilization, sections were washed twice with 1× PBS and photobleached as described before (Du et al., 2019). Following photobleaching, sections were blocked in 3% BSA (prepared in 1×PBS) for 1 h at RT. After blocking, primary antibodies were diluted in 0.5% BSA, added to the sections, and left incubating overnight at 4°C. Sections were then washed three times with 1x PBS and secondary antibodies (diluted in 0.5% BSA solution containing nuclear staining) were subsequently added and left incubating for 1 h at RT protected from light. Secondary antibodies were washed as before and coverslips were mounted on a glass slide using mounting medium. If using fluorescently labeled primary antibodies, these were added after washing the secondary antibodies and left incubating 2 h at RT. For immunofluorescence of cultured cells no acetone fixation and photobleaching were performed. For Trichrome (Masson) stainings, glass slides containing paraffin-embedded tissue sections were placed on a heating block set to 56°C for 30 min to melt the paraffin. Afterwards, sections were washed twice in xylol (5 min each) and rehydrated in decreasing alcohol concentrations (100%, 100%, 90%, 70%, 50%, 30%), for 3 min each. For Trichrome (Masson) staining, samples were left overnight in Boulin’s solution and subsequently stained following manufacturer’s instructions. Samples were mounted with a xylol-compatible mounting medium.

### Filipin and TUNEL stainings

5-8×10^4^ BALF cells were seeded in complete RPMI medium in a 24-well plate containing 10 mm sterile glass coverslips. Cells were left adhering for 3 h at 37°C. For filipin and TUNEL stainings, cells were washed with 1x PBS and subsequently fixed with 4% PFA for 30 min. For filipin staining, after washing away the fixative, cells were incubated in 100 mM glycine for 10 min at RT, and subsequently blocked with 3% BSA supplemented with 50 μg/ml filipin for 2 h at RT. Cells were washed three times with 1x PBS and immunostained as indicated above, but DRAQ5 was used as a counterstain. For TUNEL staining, manufacturer’s instructions were followed and immunofluorescence was performed after TUNEL.

### Imaging

Images of Trichrome stainings were acquired with a Zeiss Axio Observer microscope using a 20x air objective (NA 0.85). Multiple tiles, covering the entire section, with an overlap of 10% were acquired and stitched with the ZEN 3.2 software. Images of immunofluorescence of tissue sections and cultured cells were acquired using a Zeiss LSM 880 Airyscan system using a 60x oil immersion objective (NA) with a z-spacing of 500 nm. Images were acquired using the 405, 488, 561 and 640 nm laser lines. During acquisition, nuclei showing the prototypical shape of neutrophils or eosinophils were excluded.

### Image analysis

To quantify signal intensities from different markers from individual BALF cells, images were analyzed with a customized pipeline in CellProfiler. Briefly, Hoechst or DRAQ5 signals were used to segment the cells. A second primary detection step was added to create a mask of all SiglecF^+^ objects. This mask was subsequently merged onto the nuclei mask and only overlapping objects were further analyzed. A secondary object detection step was incorporated to distinguish between CD11b^+^ and CD11b^−^ cells and create a mask of alveolar macrophages. In all detection steps clumped objects were separated based on shape and signal intensities. Mean intensities for all channels for every single object were exported and used for analysis. Analysis of immunofluorescence of tissue sections was performed in QuPath. As before, Hoechst signal intensities were used to identify all objects using a radius of 2μm. For each channel, an object classifier was created to set the detection threshold based on the mean signal intensity. Subsequently, these classifiers were combined to identify alveolar macrophages. Individual cell mean intensities were exported. To measure the percentage and area of fibrosis, Trichom staining images were imported into QuPath. The total tissue area was manually annotated. A pixel classifier was trained to detect fibrotic and healthy areas. For this purpose, 5 exemplary regions of fibrotic and healthy areas were selected. Area and percentage of fibrosis were the output of this trained pixel classifier. The created classifier was applied in all the sections analyzed.

### CODEX multiplexed imaging and analysis

5 μm fresh frozen sections of the left lobe of the lung of 8-week *Ms4a3-cre^Rosa26TOMATO^* mice stimulated intranasally with PBS or β-glucan for seven days were prepared and stained following manufacturer’s instructions. Briefly, sections were retrieved from the freezer, let dry on drierite beads, and fixed in ice-cold acetone for 10 min. Following fixation, samples were rehydrated and permeabilized for 20 min with 0.2% Triton-X100. To reduce tissue autofluorescence, sections were photobleached twice for 1 h as indicated before (Du *et al*., 2019). After photobleaching, samples were washed, equilibrated for 30 min in staining buffer (Akoya Biosciences, Marlborough, MA, USA), and subsequently stained with a 17-plex CODEX antibody panel overnight at 4°C. After staining, samples were washed in staining buffer, fixed in ice-cold methanol to improve imaging quality and washed. A final fixation step with BS3 crosslinker (Sigma Aldrich) was performed. Specimens were stored in CODEX storage buffer (Akoya Biosciences) at 4°C for a maximum of one week before imaging. Antibody detection was performed in a multicycle experiment with the corresponding fluorescently-labeled reporters, following manufacturer’s instructions. Images were acquired with a Zeiss Axio Observer widefield microscope (Carl Zeiss AG, Jena, Germany) using a 20x air objective (NA 0.85) and a z-spacing of 1.5 μm. The 405, 488, 568, 647 nm fluorescent channels were used. After acquisition, images were exported using the CODEX Instrument Manager (CIM, Akoya Biosciences) and processed with the CODEX Processor v1.7 (Akoya Biosciences). Processing steps included background subtraction, stitching, shading correction, deconvolution, and cell segmentation. Cells were segmented using DAPI signals and ATPase I membrane staining to define the cell borders. Cell classification to detect alveolar macrophages and other mononuclear phagocytes was performed in CODEX MAV (Akoya Biosciences), following a similar gating scheme as the one used for flow cytometry.

### Preparation of Seq-Well arrays

Seq-Well arrays were prepared as previously described (Gierahn et al., 2017). Briefly, PDMS master mix (sylgard base, cosslinker, 10:1 ratio) was poured into a master mold and incubated for 2 h at 70°C to generate the arrays. Afterwards, the arrays were functionalized and rinsed with 100% EtOH, dried at RT and plasma treated for 7 min. After submerging in APTES and PDITC buffers, the arrays were washed in acetone and dried for 2 h at 70°C followed by incubation in 0.2% chitosan solution (pH=6.3) at 37°C for 1.5 h. After chitosan incubation, the arrays were incubated overnight in PGA buffer at RT under vacuum pressure and rotated for 3 h.

### Preparation of Seq-Well libraries

Seq-Well libraries were generated as previously described (Gierahn et al., 2017). Briefly, 110000 barcoded mRNA-capture beads in Bead Loading Buffer were loaded onto the array. Afterwards, the Bead Loading Buffer was replaced by RPMI 1640 medium (Gibco) with 10% FCS (Sigma-Aldrich) and incubated for 10 min. The medium was discarded, 20000 to 30000 BALF cells were loaded and rocked for 10 min. The loaded arrays were washed 5x with PBS, soaked in RPMI 1640 und sealed by polycarbonate membranes under mild vaccum. The sealed arrays were incubated for 37°C for 30 min in Agilent clamps (Agilent) and then incubated in a guanidinium-based lysis buffer for 20 min. After incubation in hybridization buffer, the mRNA capture beads were washed from arrays and collected. Reverse transcription was performed on the bead pellet using a Maxima Reverse Transcriptase reaction (ThermoFisher) for 30 min at room temperature with end-over-end rotation followed by 90 min incubation at 52°C with end-over-end rotation. The reaction was stopped by washing with TE buffer supplemented with 0.01% Tween-20 or 0.5% SDS. Excess primers were digested by exonuclease ExoI (New England Biolabs). Beads were counted and the reverse transcribed cDNA libraries were amplified in a PCR reaction. After PCR, 20000 – 40000 beads were pooled and cleaned using AMPure XP beads (Beckman Coulter). The library integrity was assessed using a High Sensitivity D5000 assay (Agilent) for Tapestation 4200 (Agilent).

### Sequencing

Tn5 was mixed with pre-annealed linker oligonucleotides and supplemented with glycerol, dialysis buffer and H2O. The cDNA libraries (1 ng) were tagmented with the prepared single-loaded Tn5 transposase and afterwards cleaned using MinElute PCR kit (Qiagen) following the manufacturer’s instructions. The Illumina indices (Illumina) were added to the tagmented product by PCR, which was subsequently cleaned with AMPure XP beads (Beckman Coulter). The final library quality was assessed using a High Sensitivity DNA5000 assay (Agilent) and quantified using the Qubit high-sensitivity dsDNA assay (ThermoFisher). Seq-Well libraries were equimolarly pooled and clustered at 1.4 pM concentration with 10% PhiX using High Output v2.1 chemistry (Illumina) on a NextSeq500 system (Illumina). Sequencing was performed paired-end as followed: custom Drop-Seq Read 1 primer for 21 cycles, 8 cycles for the i7 index and 61 cycles for Read 2. Single-cell data were demultiplexed using bcl2fastq2 (v2.20) (Illumina). Fastq files were loaded into a snakemake-based data pre-processing pipeline (version 0.31, available at https://github.com/Hoohm/dropSeqPipe) that relies on the Drop-seq tools provided by the McCarroll lab (Macosko et al., 2015).

### scRNAseq data analysis

Sequencing reads were mapped to the mouse reference genome mm10 using STAR alignment from the Drop-seq pipeline (v2.0.0) as previously described (Macosko, Cell, 2015). Next, we assess the quality of our libraries and excluded cells with low quality (<500 genes per cell), doublets (>3000 genes per cell), or dead cells (>10% of mitochondrial content). All genes expressed in less than 5 cells were filtered out.

Cell clustering analysis was performed using the Seurat package (v4.1.1) according to instructions (Hao et al., 2021). In brief, the expression data was log normalized with a scale factor of 10.000. After scaling, PCA was performed using the top 3000 variable genes for a linear dimensional reduction. The first 15 PCA components were used to cluster cells by the Louvain algorithm. To obtain an optimal cluster resolution, we set the resolution parameter in the FindClusters function as 0.25 to generate 5 major clusters, which were visualized after non-linear dimensional reduction with UMAP. Differentially expressed genes (DEGs) in each cluster were identified by using the default Wilcoxon rank sum test in the FindAllMarkers function, and were defined with logfc.threshold >0.25 and min.pct >0.25.

### Code and Data availability

The scRNA-seq raw reads and processed data were submitted to the NCBI GEO database accession number GSE211575. All code used for data visualization of the scRNA-seq data can be found at https://github.com/JiangyanYu/Trained_immunity_2022.

### Statistics

Statistical analysis and comparison was performed using Prism 9 (GraphPad). Data are shown as mean ± SD. Statistical significance was assessed by student’s *t*-test (unpaired) or ordinary one-way ANOVA with Tukey’s multiple comparisons test. Survival of animals is displayed in Kaplan-Meier survival curves. A p value < 0.05 was considered as statistically significant (\**P* < 0.05, \*\**P* < 0.01, \*\*\**P* < 0.001, \*\*\*\**P* < 0.0001). Mice were randomly allocated to the control or treatment groups by the investigator. Mice numbers are indicated as “n” in the figure legends.

## Author contributions

Conceptualization: H.T., D.A.B., J.H., N.K., A.S.; Formal Analysis and Investigation: H.T., D.A.B., J.H., N.K., J.S-S., K.B., J.Y., C.O.S., F.P., M.K., L.V., Y.X., M.G., L.B., A.A., S.C., K.H., G.B., D.L.W., F.G., J.R., M.B., J.L.S.; Writing:H.T., D.A.B., J.H., A.S.; Supervision:A.S.

---

**Supplement Figure 1 to Figure 1** (A-E) Quality control of the scRNA sequencing dataset of the BALF seven days after intranasal stimulation with PBS or β-glucan in wt mice. (A) Violin plot of feature gene counts per cluster. (B) Violin plot of UMI counts per cluster. (C) Violin plot of mitochondrial gene percentage per cluster. (D, E) Violin plot of *Siglecf* (D) and *Itgax* (E) RNA expression levels in the BALF seven days after β-glucan exposure by scRNA seq. (F) Percentage of contribution of condition, PBS or β-glucan, to the five clusters. (G) Flow cytometry gating strategy pre-gated on Lin^−^ (Ly6g, B220, CD19, CD3ε. Nk1.1, Ter-119) and CD45^+^ cells to define CD11b^+^ AM in the BALF after intranasal PBS or β-glucan stimulation. (H) Flow cytometric quantification of AM (CD45^+^SiglecF^+^CD64^+^CD11c^+^) numbers in the BALF in a time course from 1 to 21 days post β-glucan stimulation of wt mice (n= 5). (I) UMAP representation of intranasally stimulated lungs with PBS or β-glucan seven days after stimulation (pooled n=3 per condition). (J) Mean fluorescence Intensity (MFI) of Siglec F on indicated cell subsets in lung seven days post stimulation with β-glucan (n=4).

**Supplement Figure 2 to Figure 3** (A-C) BALF cells were harvested from wt mice seven days after intranasal exposure with PBS or 40 μl 200 μg β-glucan and subsequently restimulated *in vitro* with PBS or 10 ng/ml LPS. (A) Quantification of TNF-α protein levels by ELISA in the cell culture supernatant 24 h after restimulation with LPS (n=15). (B) Intracellular staining and flow cytometric measurement of TNF-α. Plot shows the percentage of TNF-α^+^ CD11b^+^ AM after restimulation with LPS for 6-8 h followed by intracellular staining and flow cytometric analysis (n=6). (C) Representative flow cytometry dot plots of (B) gated on CD11b^+^ AM. (D, E) Measurement of glycolytic reserve (E) and glycolytic capacity (F) in BALF cells seven days after PBS or β-glucan stimulation with Seahorse (n=6).

**Supplement Figure 3 to Figure 4** (A) Experimental setup of β-glucan-induced environmental adaptation followed by an acute infection. Intranasal PBS or β-glucan stimulation at day zero was followed by intratracheal inoculation with *Legionella pneumophila* at day seven and sacrifice two days after infection. (B) Experimental setup of β-glucan-induced environmental adaptation followed by bleomycin-induced lung fibrosis. Intranasal PBS or β-glucan stimulation at day 0 was followed by intratracheal application of *Streptomyces verticillus* bleomycin at day seven and sacrifice three or 14 days after fibrosis induction. (C, D) Disease burden and body weight of mice after pretreatment PBS or β-glucan (day 0) prior to fibrosis induction (day seven). To evaluate the disease burden (C), mice were scored for general health appearance, spontaneous behavior and body weight (n=14). (D) Body weight changes were calculated as % change based on the initial weight at day 0 (n=14). (E-G) Quantification of IL-4 (E), IL-33 (F) and TSLP (G) protein levels using Luminex Procartaplex kit 3 or 14 days after bleomycin treatment (n=7 per condition).

**Supplement Figure 4 to Figure 6** (A) Ly6c^+^ monocyte chimerism of WT/WT or WT/ApoE^flox^LysM^cre^ chimeras in BALF, blood and BM at the day of sacrifice. Bars are further subdivided to display contribution of transferred mixed bone marrow donors.

**Supplement Figure 5 to Figure 7** (A) Quantification of M-CSF protein levels in the BALF in a time course from one to 21 days post β-glucan stimulation of C57BL/6J mice (n=5). (B, C) Lungs from C57BL/6J mice were harvested 24 h after β-glucan stimulation, fixed, and frozen in OCT. 5 μm sections were prepared and stained with antibodies to identify AMs and visualize M-CSF production. Representative images are shown in (B). Arrows indicate CD11b^+^ AMs. Seven regions of the same size from samples in (B) were acquired and analyzed as explained in Materials and Methods. Plot in (C) shows the M-CSF mean signal intensities of CD11b^−^ and CD11b^+^ AMs. Each dot represents an individual cell.

