## Supplementary figures and images for "Apolipoprotein E controls Dectin-1-dependent development of monocyte-derived alveolar macrophages upon pulmonary β-glucan-induced inflammatory adaptation"

### Supplemental Figure 1

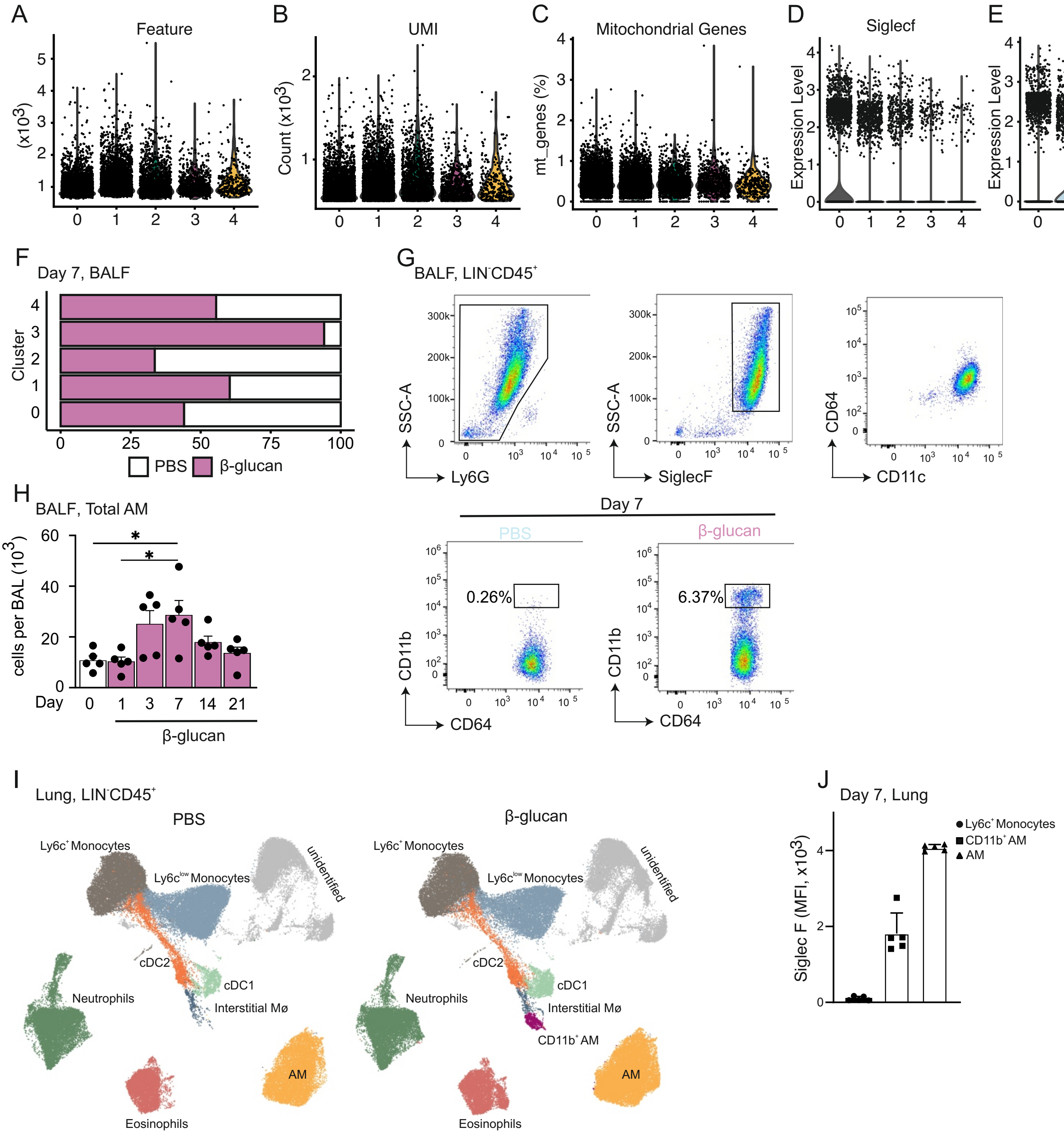

### Supplemental Figure 2

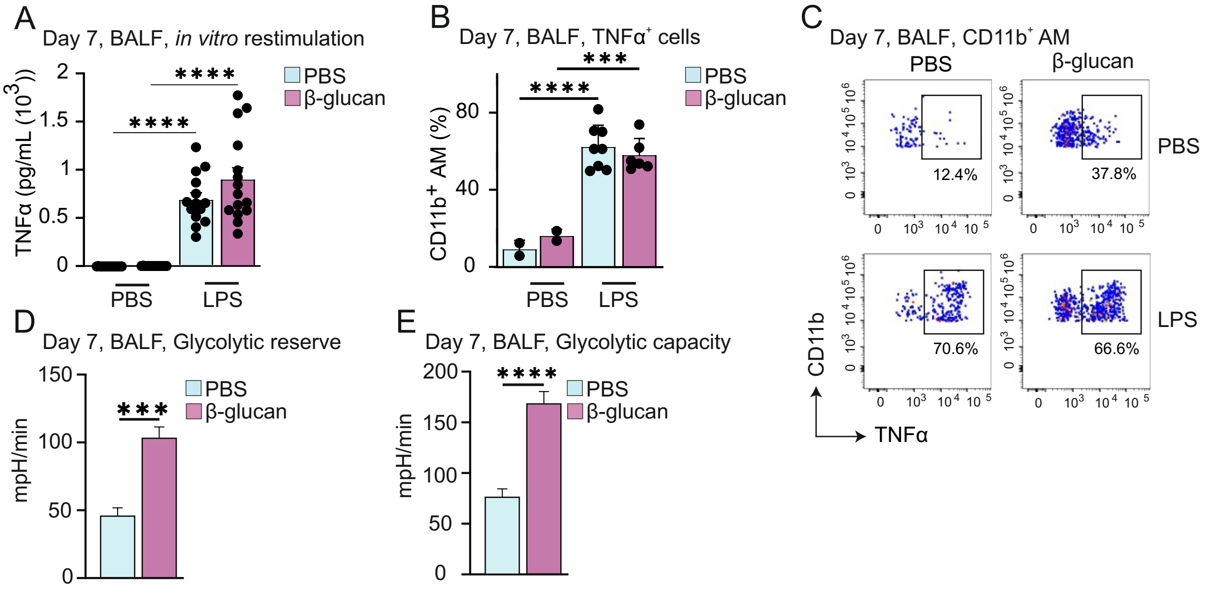

### Supplemental Figure 3

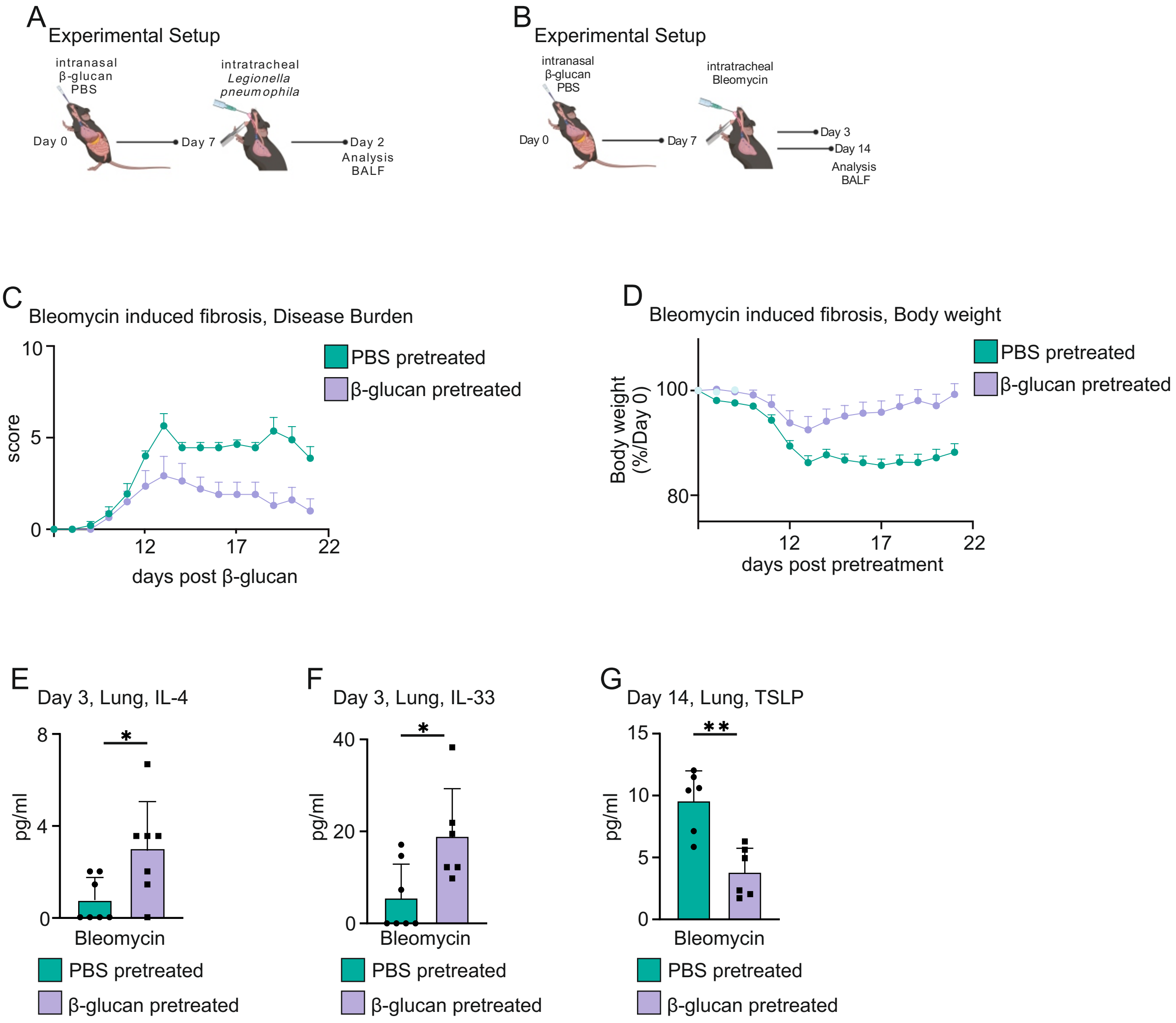

### Supplemental Figure 4

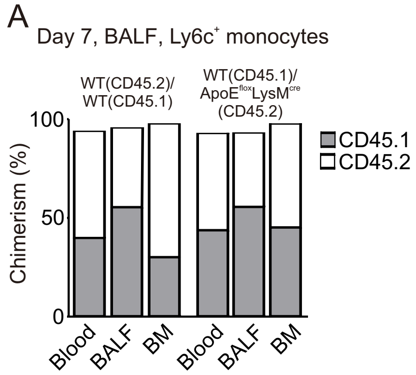

### Supplemental Figure 5

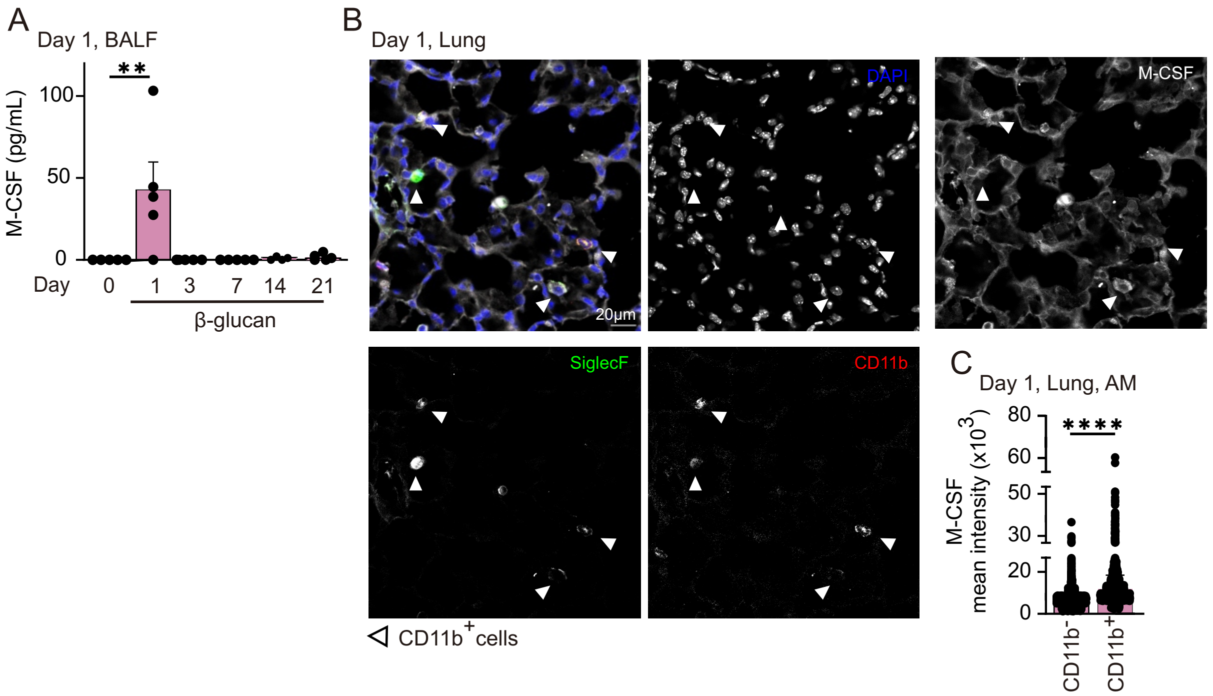
